# CHOP and IRE1α-XBP1/JNK signaling promote Newcastle Disease Virus induced apoptosis and benefit virus proliferation

**DOI:** 10.1101/300129

**Authors:** Yanrong Li, Ying Liao, Qiaona Niu, Feng Gu, Yingjie Sun, Chunchun Meng, Lei Tan, Cuiping Song, Xusheng Qiu, Chan Ding

## Abstract

Newcastle disease virus (NDV) causes severe infectious disease in poultry, and selectively kills tumor cells by inducing apoptosis. In this report, we revealed the mechanisms underlying NDV-induced apoptosis via investigation of endoplasmic reticulum (ER) stress-related unfolded protein response (UPR) in HeLa cells. We found that NDV infection induced the expression of pro-apoptotic transcription factor CHOP via PKR-eIF2α pathway. Knock down and exogenous expression studies showed that CHOP promoted cell apoptosis by down-regulation of anti-apoptotic protein BCL-2 and MCL-1, promotion of pro-apoptotic JNK and p38 signaling, and suppression of pro-survival AKT signaling. Meanwhile, CHOP facilitated NDV proliferation. Furthermore, virus infection activated IRE1α, another ER stress sensor, thereby promoting the mRNA splicing of XBP1 and resulting in the translation of transcription factor XBP1s. XBP1s entered into cell nucleus, promoted the expression of ER chaperones and components of ER associated degradation (ERAD). Exogenous expression of XBP1s helped IBV proliferation, and silence of XBP1s reduced virus proliferation. Meanwhile, exogenous expression and knock down studies demonstrated that IRE1α activated pro-apoptotic JNK signaling, promoted apoptosis and inflammation. In conclusion, our current study demonstrates that the induction of CHOP and activation of IRE1α-XBP1/JNK signaling cascades promote apoptosis and benefit NDV proliferation.

**IMPORTANCE:** It is well known that NDV kills host animal and tumor cells by inducing cell apoptosis. Although several studies investigate the apoptotic phenomena in NDV-infected tumor cells, the molecular mechanisms underlying this oncolytic virus induced apoptosis is not well understood yet. In this study, we focus on characterization of the ER stress responses in NDV-infected tumor cells, and find that virus induces apoptosis by up-regulation or activation of several unfolded protein responses (UPR) related transcription factors and signaling: such as ATF4, CHOP and XBP1s, and pro-apoptotic kinases (IRE1α, JNK, p38). Moreover, activation of these transcription factors and signaling cascades helps virus proliferation. Our study dissects the UPR induced apoptosis in NDV-infected tumor cells, and provides the evidence that UPR favors NDV proliferation.

## INTRODUCTION

The endoplasmic reticulum (ER) is a crucial intracellular organelle in eukaryotic cells. It plays important role in regulation of lipid synthesis, calcium homeostasis, and protein synthesis, translocation, folding, modification and trafficking (1). When large amounts of proteins enter the ER, unfolded or misfolded proteins accumulate in the ER lumen and induce ER stress. For survival, cell will activate several signaling pathways collectively termed as the unfolded protein response (UPR). Three transmembrane ER stress sensors have been identified, including protein kinase RNA (PKR)-like ER kinase (PERK), activating transcription factor 6 (ATF6) and Inositol-Requiring Protein 1 alpha (IRE1α) (2). In physiological condition, these sensors keep in inactive state by binding with ER chaperone immunoglobulin heavy chain-binding protein (Bip) in ER lumen. In response to excess accumulation of misfolded or unfolded proteins, Bip binds with unfolded proteins and ER sensors are released and activated. PERK is activated by homo-dimerization and auto-phosphorylation on Thr980 (2, 3), and in turn phosphorylates eukaryotic initiation factor 2α (eIF2α). Phospho-eIF2α has increased affinity to eIF2β subunit and prevents regeneration of GTP in the ternary complex eIF2-GTP-^Met^tRNAi, thus halting the ignition of protein translation (4). Although eIF2α phosphorylation leads to translation inhibition, several specific mRNAs are preferentially translated, such as activating transcription factor 4 (ATF4). ATF4 contributes to the transcription of genes important for cellular remediation and apoptosis, including growth arrest and DNA damage-inducible protein 153 (GADD153, also named CHOP) (5), a pro-apoptotic transcription factor. It is well known that CHOP is involved in cellular apoptosis by regulating the expression of BCL2, tribbles-related protein 1 (TRB3), death receptor 5 (DR5), ER Oxidoreductin-1-L-alpha (ERO1α), and DNA damage-inducible protein 34 (GADD34) (6, 7). During RNA virus infection, large amount of viral proteins are synthesized, which usually activate PERK. Meanwhile, another eIF2α kinase, PKR, binds with viral double stranded RNA (dsRNA) and is activated by auto-phosphorylation on Thr446 and Thr451 (8, 9). Both PERK and PKR are involved in phosphorylation of eIF2α on Ser51 and eliciting downstream ATF4-CHOP signaling (9, 10). Another ER stress sensor, ATF6, is released from Bip and moves to Golgi apparatus, where it is cleaved into N-terminal fragment ATF6-N and moves to nucleus as active transcription factor (11). In nucleus, ATF6-N triggers the transcription of protein chaperones, X box-binding protein 1 (XBP1), and components of ER associated degradation (ERAD), enhancing ER folding capacity and reducing misfolding proteins (12, 13). The IRE1α-XBP1 branch is the most evolutionarily conserved in Eukarya (14). Upon ER stress, IRE1α is released from Bip and undergoes homo-oligomerization, autophosphorylation, and activation. The activated IRE1α harbors the kinase activity and endoribonuclease activity(15). The endoribonuclease leads to unconventional enzymatic splicing of XBP1u mRNA into XBP1s mRNA by removing 26 nucleotide intron, and the spliced mRNA is then translated into an active transcription factor XBP1s (15). XBP1s enters into the nucleus and controls the transcription of the ER quality control genes and components of ERAD, to remove the excess misfolded/unfolded proteins in ER lumen (13, 16, 17). IRE1α also degrades ER-associated mRNAs, named as regulated IRE1α-dependent decay (RIDD), to reduce protein load in the ER (18).

If ER homeostasis cannot be restored, the UPR drives the damaged or infected cells to apoptosis (19). Apoptosis is the major type of cell death, characterized by cell shrinkage, chromosomal DNA cleavage, nuclear condensation and fragmentation, dynamic membrane blebbing, and formation of apoptosis bodies. The outside cell death ligands trigger the death receptor mediated apoptotic pathway, hallmarked by cleavage of caspase 8; the inside cell signals confer mitochondrial apoptotic pathway, hallmarked by cleavage of caspase 9. The intrinsic pathway is under control of BCL-2 protein family, which comprises at least 12 proteins, including pro-apoptotic effectors BAX and BAK, pro-apoptotic BH3-only activator proteins BID, BIM, PUMA, and NOXA, pro-apoptotic BH3-only sensitizer proteins BIK, BAD, NOXA, HRK, BNIP3, and BMF, as well as pro-survival guardian proteins BCL-2, MCL-1, BCL-XL, BCL-w, and BFL1/A1 (20). Pro-survival guardian proteins inhibit apoptosis through binding to and sequestering activators or effectors. BH3-only activators directly activate BAK or BAX. BH3-only sensitizers indirectly activate BAK or BAX through inhibiting pro-survival guardian proteins. When enough activators have been stimulated by cytotoxic stresses, BAX is released from pro-survival guardian proteins and oligomerize on the mitochondrial outer membrane, permeabilize and disrupt the membrane, resulting in release of cytochrome c and second mitochondria-derived apoptotic protein (SMAC), subsequently blocking the X-linked inhibitor of apoptotic protein (XIAP) and promoting the activation of caspase 9 on the scaffold protein apoptotic protease activating factor 1 (APAF1) (21, 22). Caspase 9 in turn cleaves and activates the effectors, such as caspase 3 (23). Under persistent ER stress, the enhanced transcription factor CHOP may promote cell apoptosis by down-regulating of anti-apoptotic protein BCL-2 expression and perturbing the cellular redox state (24). CHOP also interacts with ATF4 to induce GADD34 and recover protein synthesis (25). During prolonged ER stress, activated IRE1α interacts with TNF receptor-associated factor 2 (TRAF2), an adaptor protein, which recruits apoptosis signal-regulating kinase 1 (ASK1). The complex induces apoptosis by activation of the pro-apoptotic ASK1-c-Jun amino-terminal kinase (JNK) signaling (26). It has been demonstrated that some viruses, such as infectious bronchitis virus (IBV) and Japanese encephalitis virus (JEV), induce apoptosis via UPR in infected cells (27, 28).

Newcastle disease virus (NDV) is highly contagious avian pathogen, which belongs to the genus *Avulavirus* within the family *Paramyxoviridae* (29). Similar to other paramyxoviruses, NDV is an enveloped virus with negative-sense single-stranded RNA, which is 15186 nucleotides in length (30). The single-stranded negative RNA genome encodes six structural proteins: the hemagglutinin-neuraminidase (HN), the fusion glycoprotein (F), the matrix protein (M), the nucleoprotein (NP), the phosphoprotein (P), and the large polymerase protein (L). HN and F mediate cell surface receptor binding and membrane fusion, thereby determining virus entry into cells (31–33). M protein forms an inner protein layer below the inner leaflet of the viral membrane and plays essential role in virus assembly and budding (34). NP, P and L protein associate with the viral RNA to form the ribonucleoprotein complex (RNP), and are involved in virus genome replication (35). During the transcription of the P gene, two additional non-structural proteins, V and W, are transcribed as the result of RNA editing (36). The V protein interferes with STAT signaling and prevents interferon (IFN) stimulated genes (ISGs) expression, confers NDV the ability to evade the IFN response (37). NDV has been identified as an oncolytic virus for decades, which selectively infects and kills the human cancer tissues (38). The oncolytic activity of NDV is associated with apoptosis cascades. In NDV-infected chicken, death of chicken embryos and neurological damage in adult chicken are the consequences of the apoptosis (39). Thus, NDV infection induced apoptosis involved in the oncolytic activity and pathogenesis. It has been reported that NDV infection resulted in the loss of mitochondrial membrane potential, the release of cytochrome c, and the activation of caspase 9 (40, 41). BCL-2 and BAX modulates the NDV-induced apoptosis response (42, 43). Previously we found that NDV infection induced the expression of TNF-α and TRAIL via NF-κB pathway, thus activates caspase 8, resulting in cleavage of RIP1 and BID, thereby promoting apoptosis (44). Although both extrinsic and intrinsic apoptosis by NDV infection have been reported, the inside death signals have not been clarified yet. We have showed NDV infection induced phosphorylation of eIF2α and resulted in protein translation shut off in both cancer and chicken cells (45). However, the role of UPR in NDV-induced cell death remains largely unexplored. In this study, we focused on characterization of the UPR branches and their role in NDV-induced apoptosis. We found that NDV infection induced the expression of pro-apoptotic CHOP via PKR-eIF2α-ATF4 signaling in cancer cells. Knock down and overexpression study demonstrated that CHOP promoted NDV-induced apoptosis via reducing the level of anti-apoptotic BCL2 and MCL-1, promoting pro-apoptotic JNK and p38 signaling, and suppressing pro-survival kinase AKT. Moreover, IRE1α was activated by NDV infection, which results in the splicing of XBP1 and phosphorylation of JNK. Both IRE1α-XBP1 and IRE1α-JNK signaling play a critical role in NDV-induced apoptosis. Meanwhile, the induction/activation of CHOP, IRE1α, XBP1s, and JNK favors virus proliferation. Taken together, this study dissects the UPR branches and characterizes their roles in NDV-induced apoptosis and virus proliferation.

## MATERIALS AND METHODS

### Cells and virus

The human cervical cancer cell line (HeLa) and chicken embryo fibroblast monolayer cell line (DF-1) was purchased from ATCC (Manassas, VA, USA). These cells were cultured in Dulbecco’s modified Eagle’s medium (DMEM) (Hyclone, USA) with 4500 mg glucose supplemented with 10% fetal bovine serum (FBS, Gbico, USA) at 37°C humidified atmosphere containing 5% CO_2_.

The NDV velogenic strain Herts/33 was obtained from China Institute of Veterinary Drug Control (Beijing, China). The virus was propagated in chicken embryonate eggs and titrated on DF-1 cells by TCID_50_ assay. The virus was used for infection at multiplicity of infection (MOI) of 1 throughout this study.

### Reagents and antibodies

The IRE1α inhibitor 8-formyl-7-hydroxy-4-methylcoumarin (4μ8c) (s7272), JNK inhibitor SP600125 (s1460), and PKR/PERK inhibitor GSK2606414 (s7307) were purchased from Selleck Chemicals (USA). RNA extraction reagent Trizol^®^ and transfection reagent Lipofectamine 2000 were purchased from Invitrogen Thermo Fisher Scientific (USA). Western blotting stripping buffer (p0025) and 4’, 6’-diamidino-2-phenylindole (DAPI) (c1002) were purchased from Beyotime Biotechnology (China). SYBR Green qPCR Mix (p2092) was purchased from Dongsheng Biotech (China).

Monoclonal NDV NP antibody was raised in mice using bacterially expressed His-tagged NP as the immunogen. Antibodies against phospho-eIF2α (3398), eIF2α (5324), CHOP (2895), BCL-2 (4223), MCL-1 (5453), BCL-xL (2764), BIM (2933), PUMA (12450), BAX (5023), PARP (9542), phospho-AKT (13038), AKT (4691), phospho-ERK1/2 (4370), ERK1/2 (4695), phospho-JNK (4668), JNK (9252), phospho-p38 (4511), p38 (8690), IRE1α (3294), caspase-3 (9665) were purchased from Cell Signaling Technology (USA). Phospho-IRE1α (ab48187) and XBP1 (ab37152) was purchased from Abcam (UK). Anti-Flag and β-actin (A1978) were purchased from Sigma-Aldrich (USA). The secondary IgG conjugated with HRP, FITC, or TRITC were obtained from DAKO (Denmark).

The specific sequences of small interfering RNA (siRNA) oligos of CHOP, IRE1α, XBP1, JNK, and non-target control siRNA (sic) are shown in table 1. All siRNAs were synthesized by GenePharma Co. Ltd (Shanghai, China).

**Table 1.**
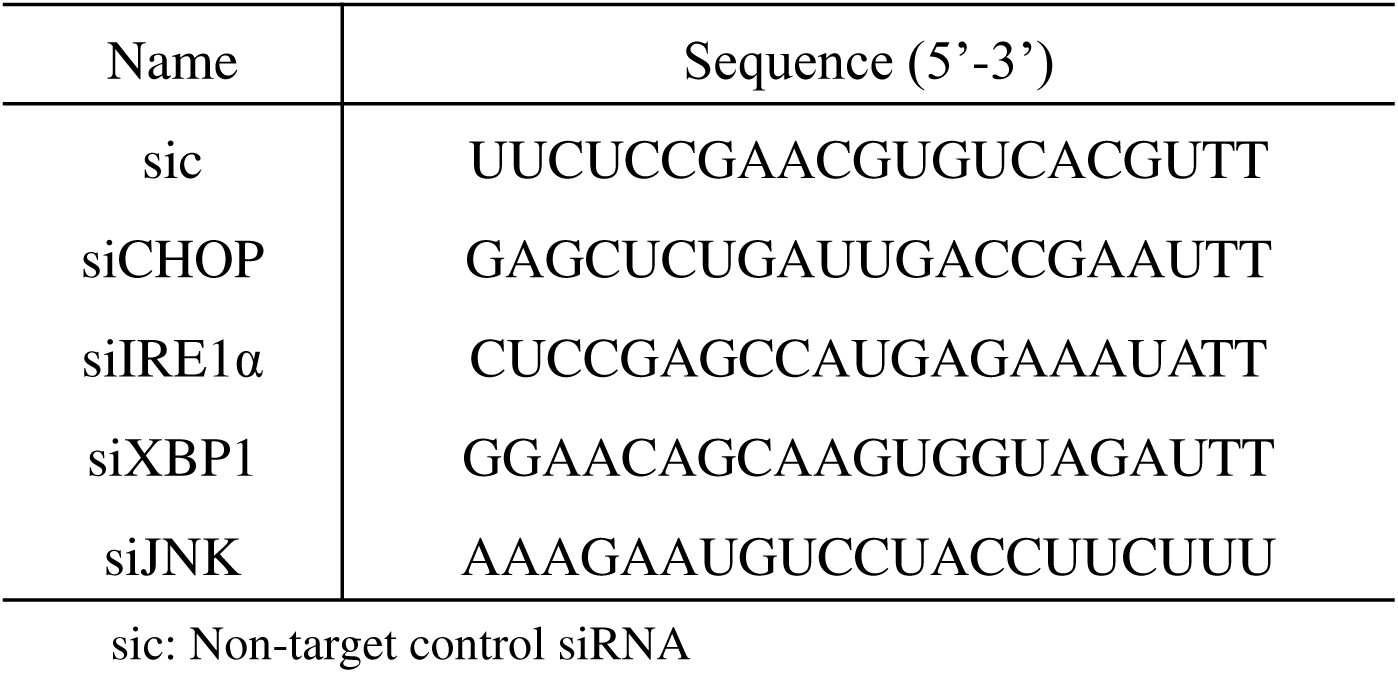
Small interfering RNA (siRNA) sequence

### Construction of plasmids

For construction of PXJ40F-CHOP plasmid, full length CHOP (NM_004083.5) was amplified by PCR from human cDNA using forward primer 5’-CCC<u>AAGCTT</u>ATGGCAGCTGAGTCATTGCCTTTC −3’ and reverse primer 5’-GGA<u>AGATCT</u>TCATGCTTGGTGCAGATTCACCATTC-3’. The restriction enzyme sites are underlined. The PCR product was digested with *Hind III* and *Bgl II* restriction enzymes, and cloned into vector PXJ40F (with a Flag tag in amino terminus). For construction of pCMV-IRE1α plasmid, full length IRE1α (GenBank: AF059198.1) was amplified by PCR from human cDNA using forward primer 5’-GCAATC<u>AAGCTT</u>ATGCCGGCCCGGCGGCTGCTGC-3’ and reverse primer 5’ -GACGTG<u>GAATTC</u>GAGGGCGTCTGGAGTCACTGGGGGC-3’. The restriction enzyme sites are underlined. The PCR product was digested with *Hind III* and *EcoR I* restriction enzymes, and cloned into vector p3xFlag-CMV-14 (with a Flag tag in carboxyl terminus). For construction of pCMV-XBP1u plasmid, full length XBP1u (NM_005080.3) was amplified by PCR from human cDNA using XBP1 forward primer 5’-GCAATC<u>AAGCTT</u>ATGGTGGTGGTGGCAGCCG-3’ and XBP1u reverse primer 5’-GACGTG<u>TCTAGA</u>GTTCATTAATGGCTTCCAGCTTGGC-3’. The restriction enzyme sites are underlined. The PCR product was digested with *Hind III* and *Xba I* restriction enzymes, and cloned into vector p3xFlag-CMV-14. For construction of pCMV-XBP1s plasmid, full length XBP1s (NM_001079539.1) was amplified by PCR from human cDNA using the forward primer 5’ -GCAATC<u>AAGCTT</u>ATGGTGGTGGTGGCAGCCG-3’ and reverse primer 5’-GACGTG<u>TCTAGA</u>GACACTAATCAGCTGGGGAAAGAG-3’. The PCR product was digested with the restriction enzyme *Pst I* to remove the XBP1u fragment, followed with *Hind III* and *Xba I* digestion, finally cloned into vector p3xFlag-CMV-14.

### Transfection of plasmids and siRNAs

HeLa cells were transfected with plasmids or siRNAs using lipofectamine 2000 reagent (Invitrogen, USA) according to the manufacture’s standard protocol. At 24 hours (h) (plasmid transfection) or 48 h (siRNA transfection) post-transfection, cells were incubated with NDV in serum free medium at 37°C for 1 h to allow the binding and entry. After that, the unbound virus was removed and cells were incubated with fresh medium (with 2% FBS). The cells and supernatant were harvested at indicated time point post-infection, and subjected to Western blotting analysis, RT-PCR, or TCID_50_ assay, respectively.

### SDS-PAGE and Western blotting analysis

Cell lysates were prepared with 2xSDS loading buffer (20 mM Tis-HCl, pH 8.0, 100 mM Dithiothreitol, 2% SDS, 20% Glycerol and 0.016% Bromphenol blue) and denatured at 100°C for 5 min. The whole cell lysates were separated by SDS-PAGE and transferred onto nitrocellulose membranes (Sigma-Aldrich, USA). The membranes were blocked with 5% fat free milk in Tris-buffered saline with 0.05% Tween 20 (TBST) for 1 h, and incubated with the primary antibodies (1:1000 in dilution) overnight at 4°C, then washed thrice with TBST. The membranes were then incubated with secondary antibody (1:1000 in dilution) for 1 h at room temperature and washed thrice with TBST. The protein bands were detected by enhanced chemiluminescence (ECL) detection system (Share-Bio, Shanghai, China) and exposed to Automatic chemiluminescence image analysis system (Tanon, 5200, China). After the detection, membranes were washed for 5 minutes (min) with TBST, followed by rinsing with Western blotting stripping buffer for 20 min. Then, the membranes were rinsed with TBST and blocked with 5% fat free milk in TBST before re-probing with other antibodies.

The intensities of target bands were quantified using Image J program (NIH, USA).

### Immunofluorescence

HeLa cells were grown on 4-well chamber slides and infected with NDV. At 16 hours post-infection (h.p.i.), cells were fixed with 4% paraformaldehyde for 15 min, permeabilized with 0.5% Triton X-100 for 10 min, and blocked with 3% BSA for 30 min. The cells were incubated with antibody against CHOP or XBP1, and NDV NP (1:200 dilution, 5% BSA) for 1 h, respectively, followed by staining with secondary antibody conjugating with FITC or TRITC (1:200 dilution, 5% BSA) for another 1 h. Finally, cell nuclei were stained with 0.1 μg/ml of DAPI for 10 min and rinsed with PBS. The specimen was mounted with fluorescent mounting medium (DAKO) containing 15 mM NaN3. Images were collected with a LSM880 confocal laser-scanning microscope (Zeiss, German).

### Semi-quantitative real time RT-PCR

Total RNA was extracted using TRIzol^®^ Reagent (Invitrogen, USA) according to the manufacturer’s instructions. Briefly, cells were lysed with TRIzol and the lysates were mixed with one-fifth volume of chloroform. After centrifugation at 12000×g at 4°C for 15 min, the aqueous phase was mixed with an equal volume of isopropanol. RNA was pelleted by centrifugation at 12000×g at 4°C for 20 min, washed with 70% ethanol twice, and dissolved in RNase-free H_2_O. The concentration of the RNA was measured using a NanoDrop 2000 spectrophotometer (Thermo Fisher Scientific, USA).

cDNA was reversed transcribed from total RNA using expand reverse transcriptase (Roche, USA) and oligo-dT primer. Equal volume of cDNA was PCR-amplified using SYBR Green qPCR Mix in a CFX96TM real-time PCR system (Bio-Rad, USA). Primers used for amplify β-actin, NP, IRE1, XBP1u, XBP1s, P58^IPK^, ERdj4, EDEM1, IFN-β, TNF-α, IL-6 and IL-8 were listed in table 2. The mRNA levels of specific genes were calculated using β-actin as an internal reference and normalized to mock sample. All assays were performed in three replicates.

**Table 2.**
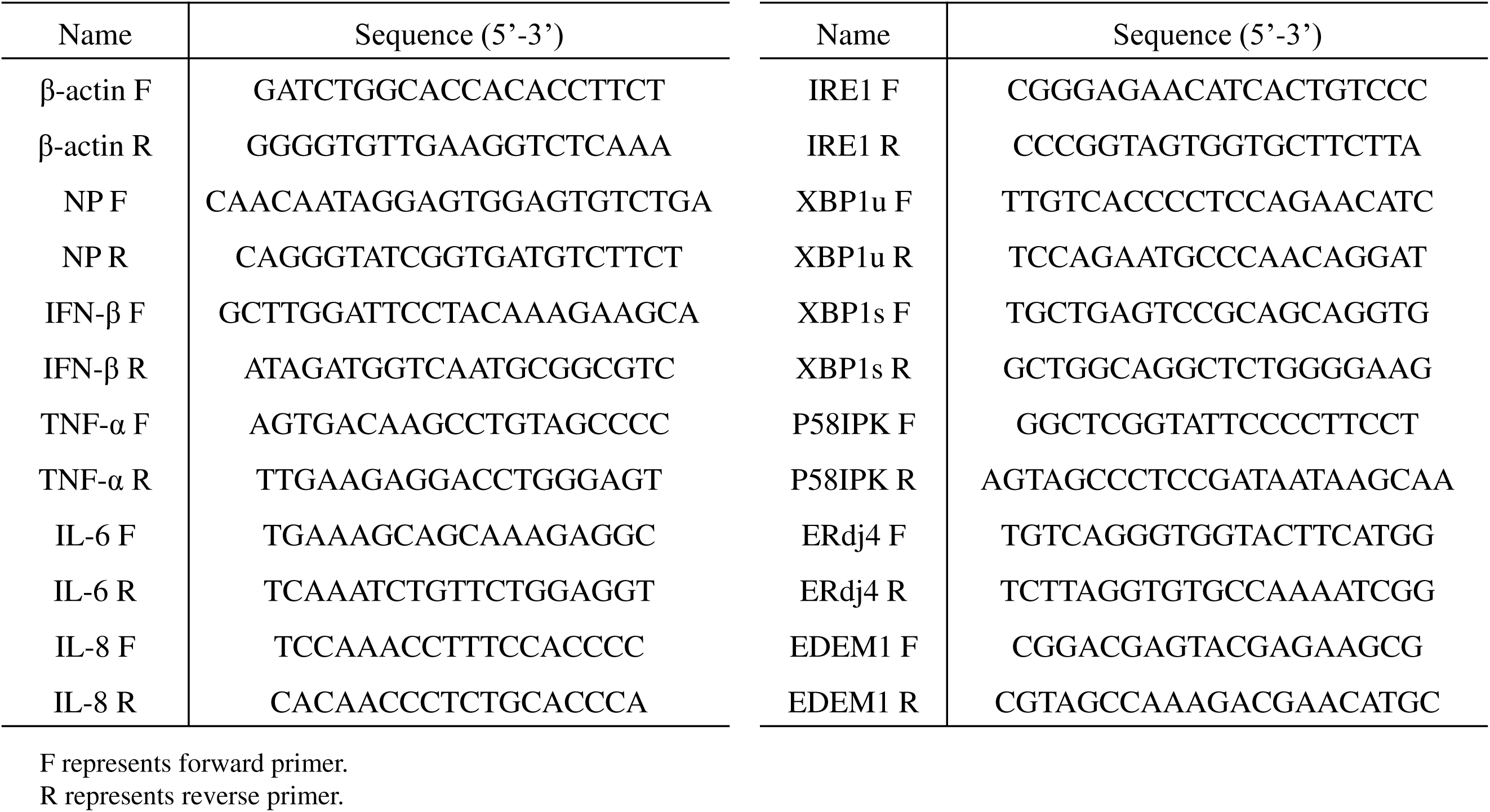
Primer sequences used for semi-quantitative real time RT-PCR

The XBP1 splicing was checked by RT-PCR using forward primer 5’-CCAAGGGGAATGAAGTGAGGC-3’ and reverse primer 5’-AGAGTTCATTAAT GGCTTCCAG-3’, which produces unspliced XBP1 of 335 bp and spliced XBP1 of 309 bp. The PCR products were digested with the restriction enzyme *Pst I*, cleaving XBP1u into 72 bp and 263 bp. The digestion products were resolved on 2.5% agarose gel to separate unspliced and spliced XBP1.

### Tissue Culture Infectious Dose 50 (TCID_50_) Assay

Virus yield in culture medium of NDV-infected cells was determined by measuring TCID_50_ in DF-1 cells. In brief, DF-1 cells were seeded in 96-well plates at a density of 2.0×10^4^ cells per well. After 24 h, cells were infected with virus sample, which was serially diluted in 10-fold using serum free medium. The virus and cells were incubated at 37°C for 4 days. The cytopathic effect of cells was observed using light microscopy. TCID_50_ was calculated by the Reed-Muench method.

### Statistical analysis

The statistical analysis was performed with Graphpad Prism5 software (USA). The data were expressed as means ± standard deviation (SD) at least three independent experiments. Significance was determined with the one-way analysis of variance (ANOVA). P values < 0.05 were deemed statistically significant.

## RESULTS

### NDV infection induces the expression of pro-apoptotic transcription factor CHOP in time-dependent manner

Our previous studies have reported that the ER stress response branch PERK/PKR-eIF2α-ATF4-GADD34 is activated in tumor and chicken cells infected with NDV (45, 46). As ER stress is a dynamic process, whether ER stress is pro-survival or pro-apoptotic dependents mostly depend on the duration and extent of the ER stress (47). Under prolonged ER stress, the preferentially translation of ATF4 usually promotes the expression of pro-apoptotic transcription factor CHOP (48, 49). Therefore, we measured the levels of this key ER stress pro-apoptotic marker CHOP after exposure with NDV. Human cervical cancer cells HeLa were either infected with NDV at MOI of 1 or mock-infected, followed by Western blotting analysis. As shown in Fig. 1A, the expression of CHOP was almost undetectable in mock-infected cells; however, it was elevated and accumulated by 10.7 to 24.9-fold during NDV infection at 12-24 h.p.i.. To check whether CHOP enters into nucleus as active transcription factor, immunofluorescence assay was performed at 16 h.p.i.. Fig. 1B showed that CHOP was barely detectable in mock-infected cells; however, in NDV-infected cells, CHOP signal was intensified and mainly localized within the nucleus. The induction of CHOP by NDV infection is also demonstrated in lung cancer cells A549 (data not shown). Thus, NDV infection greatly induces the expression of CHOP in time dependent manner and promotes its nuclear translocation. The persistent exposure to the NDV infection results in pro-apoptotic transcription factor expression and may promote apoptotic death.

**Figure 1.**
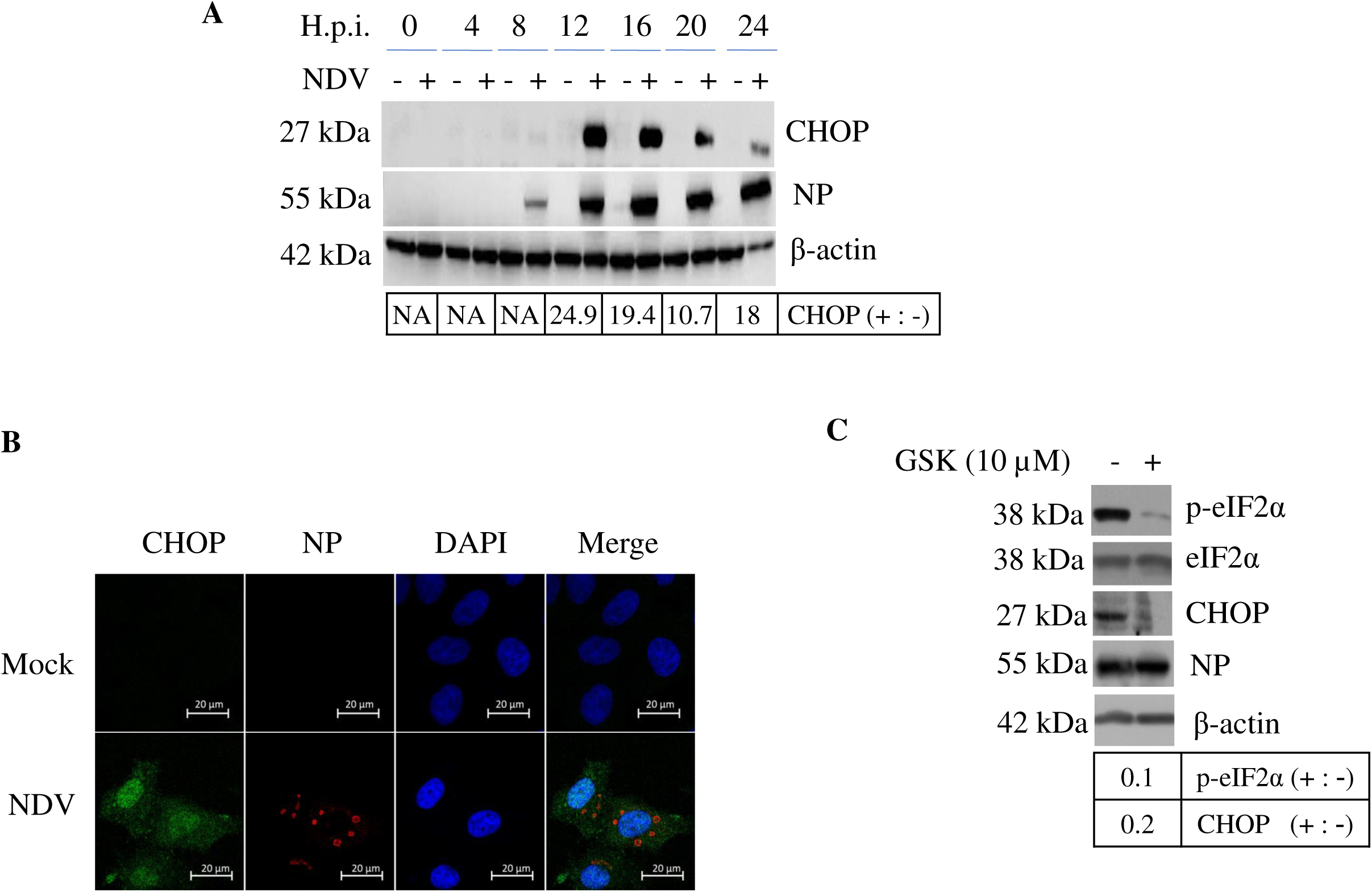
NDV infection induces the expression of transcription factor CHOP. (A) Induction of CHOP by NDV infection. HeLa cells were infected with NDV and harvested at 0, 4, 8, 12, 16, 20, and 24 h.p.i.. The cell lysates were analyzed by Western blotting with antibodies against CHOP and NDV NP protein. β-actin was detected as a loading control. The intensity of CHOP band was determined by Image J software and normalized to β-actin, respectively, and shown as fold change of NDV (+ : -). (B) Nuclear translocation of CHOP during NDV infection. HeLa cells were infected with NDV and subjected to immunofluorescence at 16 h.p.i., using antibodies against CHOP and NP. The signal of CHOP and viral protein NP were observed under confocal microscope. (C) CHOP is induced by PKR-eIF2α signaling. HeLa cells were infected with NDV and treated with 10 μM of GSK2606414 (GSK), and harvested at 16 h.p.i.. DMSO treatment was included as control in a parallel experiment. The phospho-eIF2α, CHOP, NP, and β-actin were analyzed with Western blotting. The intensity of phospho-eIF2α and CHOP band was determined, normalized to eIF2α or β-actin respectively, and shown as fold change of GSK (+ : -).

During our previous study, we found that PERK was cleaved and PKR was responsible for the eIF2α phosphorylation and GADD34 induction during NDV infection (45). To clarify whether PERK-eIF2α or PKR-eIF2α signaling is involved in NDV-induced CHOP expression, HeLa cells were treated with GSK2606414, an inhibitor blocks PERK activity at low dose (IC_50_: 0.4 nM) and blocks PKR activity at high dose (IC_50_: 696 nM) (50). In our previous report, we have demonstrated that GSK2606414 did not decrease the phosphorylation level of eIF2α at the low dose of inhibiting PERK(45). Thus, PKR may be responsible for phosphorylation of eIF2α and CHOP induction. Thus, we treated NDV-infected HeLa cells with 10 μM of GSK2606414, a dose suppressing PKR activity. Western blotting results showed this dose of inhibitor greatly reduced phospho-eIF2α by 0.1-fold and decreased CHOP by 0.2-fold, compared to DMSO-treated group (Fig. 1C). Therefore, PKR may contribute to the activation eIF2α-ATF4-CHOP pathway during NDV infection.

### CHOP promotes NDV-induced apoptosis by reducing the level of anti-apoptotic protein BCL-2 and MCL-1

We have reported that NDV infection induces both intrinsic and extrinsic apoptosis, and the extrinsic apoptosis is mediated by induction of death ligands, such as TNF-α and TRAIL (44). Whether CHOP is involved in NDV-induced intrinsic apoptosis? It has been known that CHOP promotes mitochondria mediated apoptosis via down-regulation of the pro-survival BCL-2 family (24, 51). Thus, it will be interesting to check whether the expression level of BCL-2 family is regulated by NDV infection. Western blotting analysis showed that pro-survival BCL-2 and MCL-1 were gradually decreased by 0 to 0.5-fold from 16 to 24 h.p.i in NDV-infected HeLa cells (Fig. 2A). However, BCL-xL remained relatively stable along the infection time course (Fig. 2A). Moreover, the pro-apoptotic BH3 only proteins BIM and PUMA, the pore forming protein BAX and BAK, kept in steady level (Fig. 2A). The decrease of BCL-2 and MCL-1 implies that more BAX and BAK are released and form pores in mitochondria outer membranes, initiating apoptosis. To investigate the role of CHOP in regulation of BCL-2 and MCL-1 level during NDV infection, we used siRNA (siCHOP) to specifically knock down CHOP in HeLa cells. Cells were transfected with siCHOP or non-targeting control siRNA (sic), followed with NDV infection at 36 h post-transfection, and subjected to Western blot analysis at 16 h.p.i.. As expected, compared to sic transfected control group, siCHOP efficiently knocked down the NDV-induced expression of CHOP (Fig. 2B). As expected, knock down of CHOP increased the level of MCL-1 and BCL-2 by 2.6-fold and 1.1-fold, respectively (Fig. 2B). Cleavage of poly ADP-ribose polymerase (PARP), a substrate of caspase-3, from the 116-kDa full length protein (PARP-FL) to an 85-kDa inactive polypeptide (PARP-C), was used here as a major biochemical marker of apoptosis. As shown in Fig. 2B, a significant amount of the PARP cleavage product was detected in NDV-infected control group; in contrast, less PARP cleavage (0.6-fold) was observed in NDV-infected CHOP knock down cells. Above results substantiates the hypothesis that CHOP plays a pro-apoptotic role in NDV-infected cells, probably through regulation of MCL-1 and BCL-2 level. Surprisingly, viral NP expression level was reduced by 0.4-fold in CHOP knock down cells compared to that in control cells (Fig. 2B). Accordingly, the release of virus progeny was greatly reduced, as determined by TCID_50_ assay (Fig. 2C). The experiment was performed multiple times and reproducible. Above results demonstrate that virus proliferation is moderately suppressed in CHOP knock down cells.

**Figure 2.**
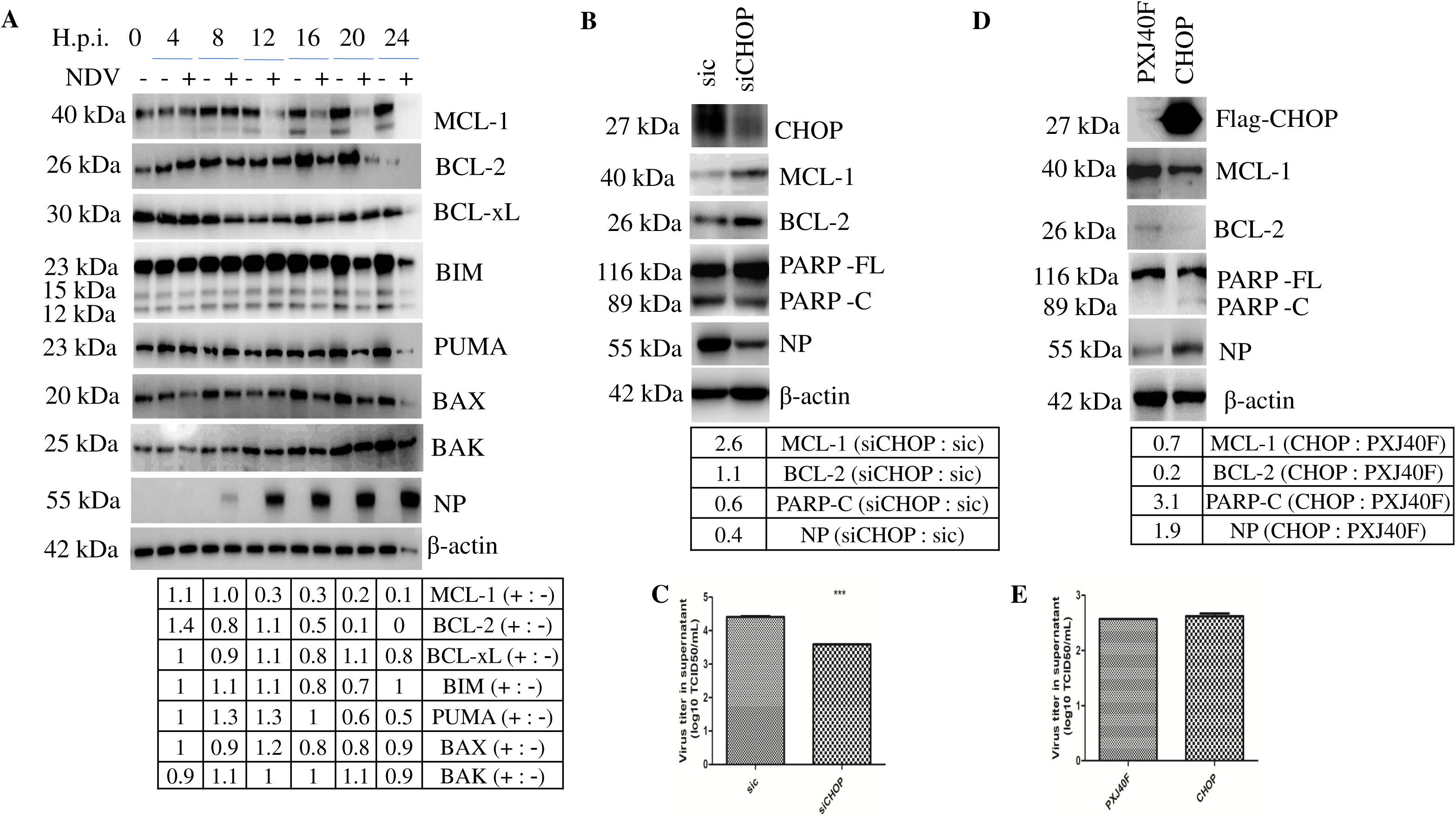
CHOP promotes apoptosis by down-regulation of anti-apoptotic protein BCL-2 and MCL-1 during NDV infection. (A) Down-regulation of BCL-2 and MCL-1 during NDV infection. HeLa cells were infected with NDV and harvested at 0, 4, 8, 12, 16, 20, and 24 h.p.i.. Western blotting analysis was performed to detect BCL-2, MCL-1, BCL-xL, BIM, PUMA, BAX, BAK, NDV NP, and β-actin. The intensity of indicated protein bands was determined, normalized to β-actin, and shown as fold change of NDV (+ : -). (B-C) Knock down of CHOP by siRNA recovers the level of BCL-2 and MCL-1, reduces apoptosis, and suppresses virus proliferation. HeLa cells were transfected with siCHOP or sic, followed with NDV infection. Cell lysates were prepared at 16 h.p.i. and analyzed with Western blotting using antibodies against CHOP, MCL-1, BCL-2, PARP, NP, and β-actin. The intensity of indicated protein bands was determined, normalized to β-actin respectively, and shown as fold change of siCHOP: sic (B). Meanwhile, the virus progeny in culture medium was titrated with TCID_50_ assay (C). (D-E) Overexpression of CHOP down-regulates MCL-1 and BCL-2, promotes apoptosis, and facilitates viral proliferation. HeLa cells were transfected with PXJ40F-CHOP or PXJ40F, followed with NDV infection. Cells were harvested at 16 h.p.i., and analyzed with Western blotting using antibodies against CHOP, MCL-1, BCL-2, PARP, NP, and β-actin. The intensity of indicated protein bands was determined, normalized to β-actin respectively, and shown as fold change of CHOP: PXJ40F (D). The virus progeny in culture medium was titrated with TCID_50_ assay (E).

To confirm above observation, we next adopted the transient overexpression approach. A plasmid encoding the full-length human CHOP with Flag tag at the N terminus was constructed. HeLa cells were transfected with the construct or vector control (PXJ40F) for 24 h before being infected with NDV for 16 h. As shown in Fig. 2D, the successful expression of Flag-CHOP was detected with Western blotting using antibody against Flag tag. Compared with that in vector control cells, overexpression of CHOP reduced the level of MCL-1 by 0.7-fold and BCL-2 by 0.2-fold, respectively. As expected, overexpression of CHOP promoted PARP cleavage by 3.1-fold. Meanwhile, the expression of viral protein NP was increased by 1.9-fold in CHOP transfected cells (Fig. 2D). Furthermore, the virus yield in culture medium was also increased (Fig. 2E). The experiment was performed multiple times and reproducible. Taken together, these data further demonstrate that CHOP promotes apoptosis via down-regulation of BCL-2 and MCL-1, and helps NDV proliferation.

### CHOP promotes apoptosis by regulation of AKT and JNK/p38 signaling cascades

MAPK cascades play a critical role in regulation of cell growth, differentiation, and control of cellular responses to cytokines and stress (52, 53). ERK1/2 is activated by growth and neurotrophic factors (54–56); JNK and p38 MAPK are activated by inflammatory cytokines and by a wide variety of cellular stresses (57, 58). AKT plays a critical role in promoting cell survival by inhibiting apoptosis (59), through phosphorylation and inactivation of several targets, including Bad (60), forkhead transcription factors (61), c-Raf (62), and caspase 9 (63). To check whether MAPK and AKT pathways are involved in NDV-induced apoptosis, the kinetic activation of these kinases during NDV infection was examined by Western blotting analysis. As shown in Fig. 3A, AKT was phosphorylated from 4 to 24 h.p.i. in both mock- and NDV-infected cells, compared to that at 0 h.p.i.. This might be due to stimulation of AKT signaling by removing serum during infection. It was noted that the level of phospho-AKT in NDV-infected cells was higher by 2.8- to 10.3-fold than that in mock-infected cells at 16-24 h.p.i., suggesting the virus infection moderately stimulates AKT signaling at late infection stage. The level of phospho-ERK1/2 was also increased from 4 to 24 h.p.i. in both mock- and NDV-infected cells, compared to that at 0 h.p.i.. Also, the level of phospho-ERK1/2 in NDV-infected cells was higher than that in mock-infected cells at 12-24 h.p.i., indicating the virus infection stimulates ERK1/2 signaling at late infection stage. A gradual increase in phospho-JNK (34.5- to 120-fold) and phospho-p38 (1.7- to 23.9-fold) at 12-24 h.p.i. were detected in NDV-infected cells, both of which were almost undetectable in mock-infected cells. Above results reveals that NDV infection moderately activates pro-survival AKT and ERK1/2, and greatly stimulates pro-apoptotic JNK and p38 signaling at late infection stage.

**Figure 3.**
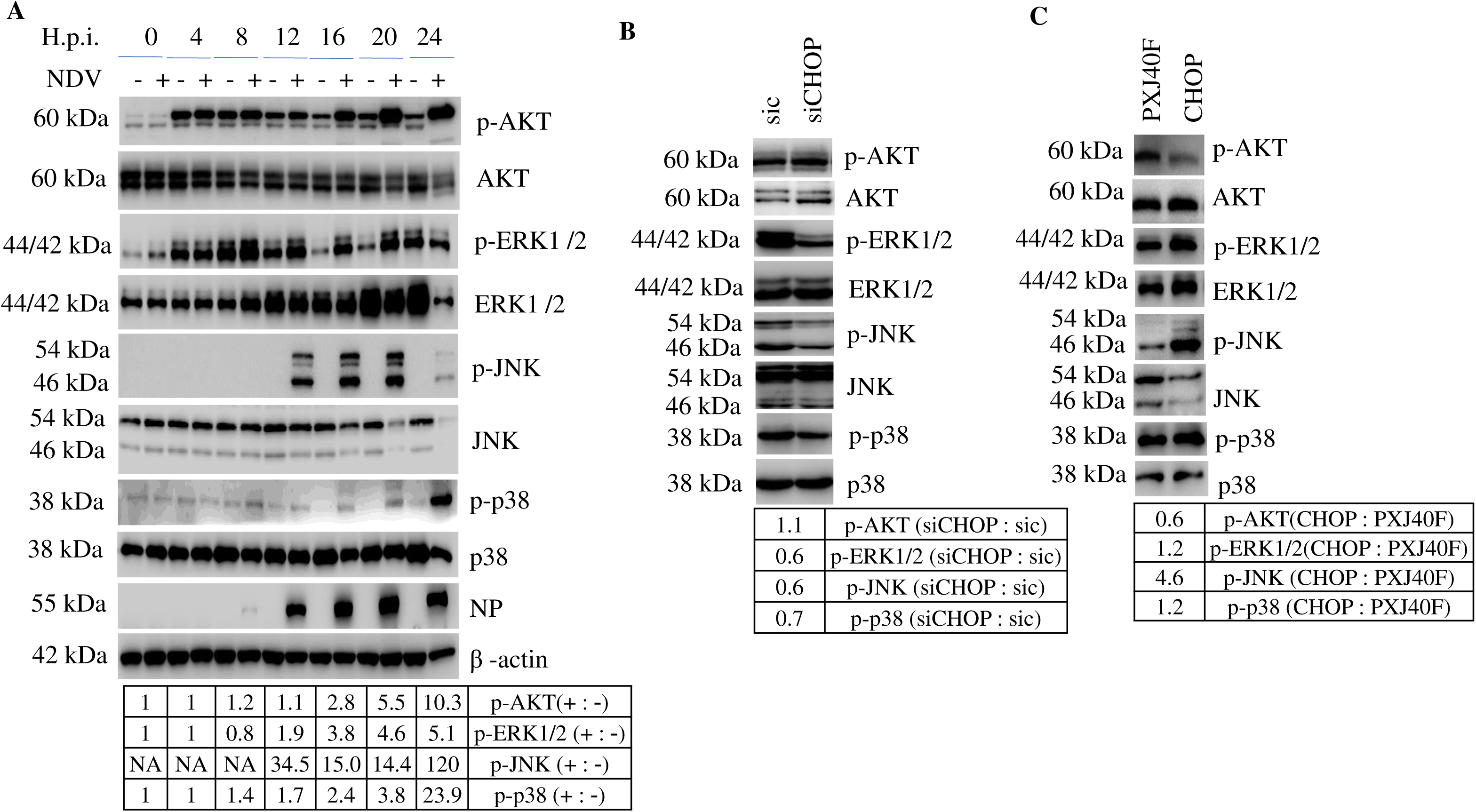
CHOP promotes apoptosis by suppression of the AKT signaling cascade and promotion of JNK/p38 signaling cascades. (A) Activation of AKT, ERK1/2, JNK, and p38 signaling cascades during NDV infection. HeLa cells were infected with NDV and harvested at indicted time points. Western blotting analysis was performed using antibodies against phospho-AKT, AKT, phospho-ERK1/2, ERK1/2, phospho-JNK, JNK, phospho-p38, p38, NP, and β-actin. The intensity of phospho-AKT, phospho-ERK1/2, phospho-JNK, and phospho-p38 bands was determined, normalized to respective total protein, and shown as fold change of NDV (+ : -). (B) Depletion of CHOP by siRNA knock down slightly increases the AKT signaling cascades and greatly suppresses MAPK signaling cascades. HeLa cells were transfected with siCHOP or sic, followed with NDV infection. The cell lysates were prepared at 16 h.p.i and analyzed with Western blotting. The intensity of phospho-AKT, phospho-ERK1/2, phospho-JNK, and phospho-p38 bands was normalized to respective total protein and shown as fold change of siCHOP: sic. (C) Overexpression of CHOP suppresses AKT signaling cascades and stimulates MAPK signaling cascades. HeLa cells were transfected with PXJ40F-CHOP or PXJ40F, followed with NDV infection. The cell lysates were prepared at 16 h.p.i. and analyzed with Western blotting. The intensity of phospho-AKT, phospho-ERK1/2, phospho-JNK, and phospho-p38 bands was normalized to respective total protein and shown as fold change of CHOP: PXJ40F.

To study whether CHOP is involved in regulation of above signaling cascades, CHOP was either knocked down or overexpressed in HeLa cells, followed by NDV infection. As shown in Fig. 3B and 3C, knock down of CHOP slightly increased the phosphorylation of AKT by 1.1-fold; however, overexpression of CHOP greatly reduced the level of phospho-AKT by 0.6-fold, suggesting that CHOP inhibits the pro-survival AKT signaling. The levels of phospho-ERK1/2, phospho-JNK, and phospho-p38 were remarkably reduced by 0.6 to 0.7-fold in knock down cells (Fig. 3B); however, overexpression of CHOP augmented the NDV-induced activation of all the three MAPKs by 1.2 to 4.6-fold (Fig. 3C). From these evidences, we speculates that augmentation of three MAPK pathways and inhibition of pro-survival AKT signaling by CHOP may play a functional role in promoting NDV-induced apoptosis during NDV infection.

### Activation of IRE1α promotes NDV-induced apoptosis and facilitates viral replication

IRE1α belongs to the evolutionarily oldest branch of the UPR in mammals. During ER stress, the kinase and RNase domains of IRE1α are activated cooperatively (64). IRE1α signaling pathway has shown to be involved in apoptotic cell death under prolonged/severe ER stress (26, 65). To check whether NDV infection activates the IRE1α signaling, phosphorylation of IRE1α during NDV infection was examined. As shown in Fig. 4A, NDV infection greatly stimulated the phosphorylation of IRE1α by 2.7- to 24.4-fold from 12 to 24 h.p.i., compared to that in mock-infected group. To access the role of IRE1α in NDV-induced apoptosis, we manipulated this protein expression by siRNA knock down. HeLa cells were transfected with siIRE1α or sic for 36 h, followed with NDV infection for 16 h. The knock down efficiency was determined by Western blotting. As shown in Fig. 4B, the expression of IRE1α was successfully knocked down by siRNA, as evidenced by undetectable level of phospho-IRE1α and total IRE1α. This knock down led to less cleavage of apoptosis marker protein caspase 3 (0.4-fold) and PARP (0.25-fold), compared to those in sic-transfected cells. These results demonstrate that IRE1α plays a crucial role in NDV-induced apoptosis. Meanwhile, viral protein NP expression was significant suppressed by 0.2-fold in IRE1α knock down cells (Fig. 4B). In consistence, in the absence of IRE1α, NP mRNA transcription was decreased by 0.25-fold, as determined by semi-quantitative real time RT-PCR (Fig. 4C); virus particles released in culture medium were also reduced, as confirmed by TCID_50_ assay (Fig. 4D). Taken together, these results reveal that IRE1α is essential for NDV proliferation and promotes the infected cells to apoptosis.

**Figure 4.**
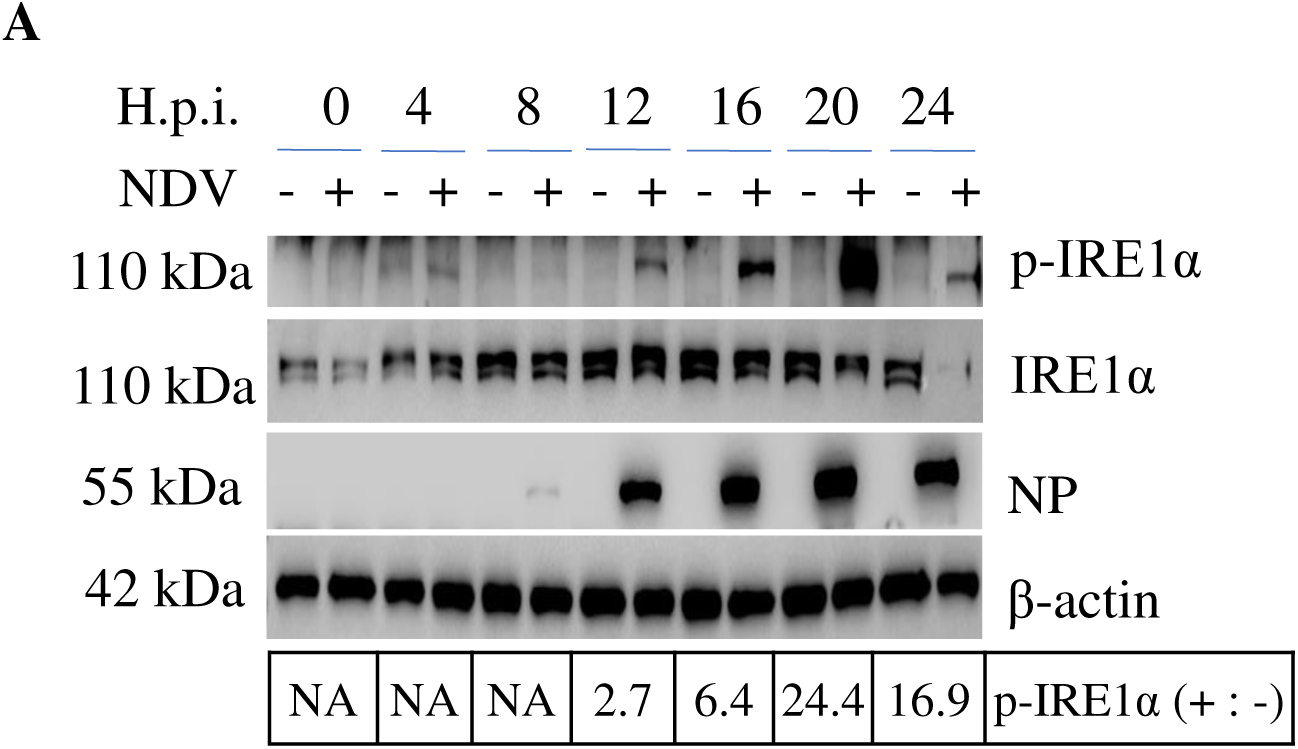
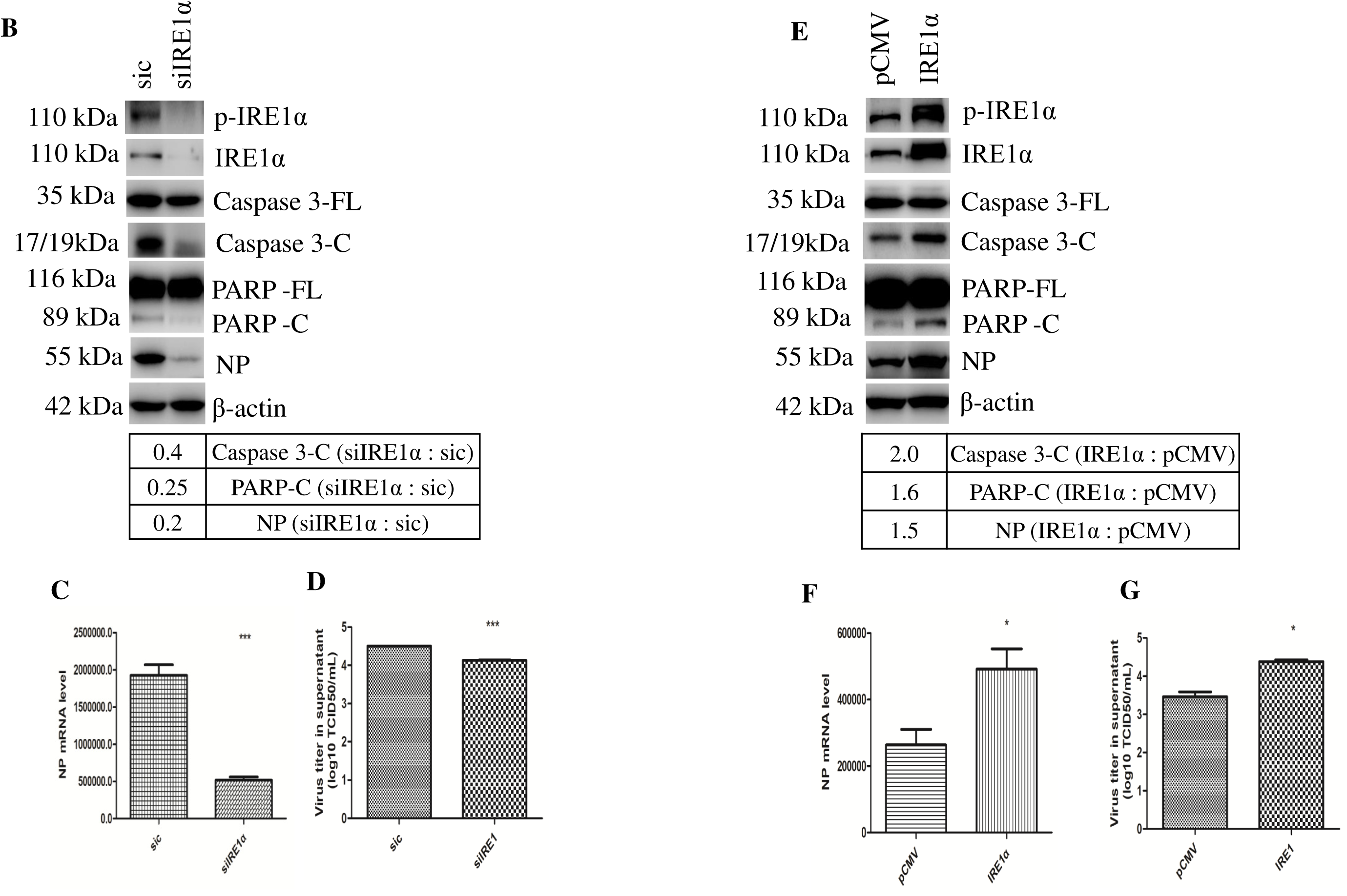
IRE1α promotes NDV-induced apoptosis and benefits NDV proliferation. (A) Activation of IRE1α in NDV-infected cells. HeLa cells were infected with NDV and harvested at indicated time points. Cell lysates were analyzed with Western blotting using antibodies against phospho-IRE1α, IRE1α, NP, and β-actin. The intensity of phospho-IRE1α bands was normalized to total IRE1α and shown as fold change of NDV (+ : -). (B-D) Knockdown of IRE1α reduces apoptosis and virus proliferation. HeLa cells were transfected with siIRE1α or sic, followed with NDV infection. The cell lysates were prepared at 16 h.p.i. and analyzed with Western blotting to detect phospho-IRE1α, IRE1α, caspase-3, PARP, NP, and β-actin. The intensity of caspase-3-C, PARPHOSPHO-C, and NP bands was compared to caspase-3-FL, PARPHOSPHO-FL, or β-actin, and shown as fold change of siIRE1α : sic (B). Meanwhile, semi-quantitative real time RT-PCR was performed to detect NP mRNA (C), and the virus titer in culture medium was titrated with TCID_50_ assay (D). (E-G) Overexpression of IRE1α augments NDV-induced apoptosis and promotes virus proliferation. HeLa cells were transfected with pCMV-IRE1α or pCMV, followed with NDV infection. The cell lysates were prepared at 16 h.p.i. and analyzed with Western blotting to detect phospho-IRE1α, IRE1α, caspase-3, PARP, NP, and β-actin. The intensity of caspase-3-C, PARPHOSPHO-C, and NP bands was compared to caspase-3-FL, PARPHOSPHO-FL, or β-actin, and shown as fold change of IRE1α: pCMV (E). Meanwhile, semi-quantitative real time RT-PCR was performed to detect NP mRNA (F), and the virus progeny in culture medium was titrated with TCID_50_ assay (G).

To validate above conclusion, we further analyzed the apoptosis and virus proliferation in IRE1α overexpressing cells. A plasmid encoding full length human IRE1α with Flag at C-terminus was constructed. HeLa cells were transfected with the construct or vector before IBV infection. As shown in Fig. 4E, compared with vector pCMV transfection, transfection of IRE1α construct resulted in higher level of phospho-IRE1α and IRE1α. This resulted in more cleavage of apoptosis marker protein caspase 3 (2.0-fold) and PARP (1.6-fold), compared to those in sic-transfected cells. Furthermore, compared with that in control cells, 1.5-fold of viral NP protein production was observed in IRE1α overexpressing cells (Fig. 4E); similarly, NP mRNA was increased by 2-fold (Fig. 4F), more virus particles were released into culture medium (Fig. 4G). Altogether, above results confirm that activation of IRE1α promotes NDV-induced apoptosis and is necessary for efficient virus replication. Why IRE1α is so important in cell death and NDV proliferation? The underlying mechanisms need further exploration.

### XBP1 is spliced by IRE1α and promotes the expression of ER chaperones

The activated IRE1α catalyzes the splicing of XBP1 mRNA by removing a 26 nucleotide intron, producing XBP1s mRNA, which is translated into 55 kDa XBP1s as active transcription factor (12, 16). We next examined the splicing of XBP1 by IRE1α during NDV infection by Western blot and RT-PCR. As shown in Fig. 5B, the 55 kDa XBP1s protein was observed at 12 h.p.i. and gradually increased at 16-24 h.p.i., while the 40 kDa unsplicing isoform XBP1u was decreased along infection time course (Fig. 5A). Consistent with above result, RT-PCR analysis detected the increase of XBP1s mRNA by 1.5- to 8.9-fold at 12-20 h.p.i. (Fig. 5B). To access whether XBP1s really enters into nucleus as transcription factor, immunofluorescence was performed at 16 h.p.i.. The image in Fig. 5C revealed that XBP1 was diffused in cytoplasm in mocked-infected cells, and entered into nucleus during NDV infection. Above results clearly demonstrate that NDV infection promotes XBP1 mRNA splicing and produces XBP1s protein as an active transcription factor.

**Figure 5.**
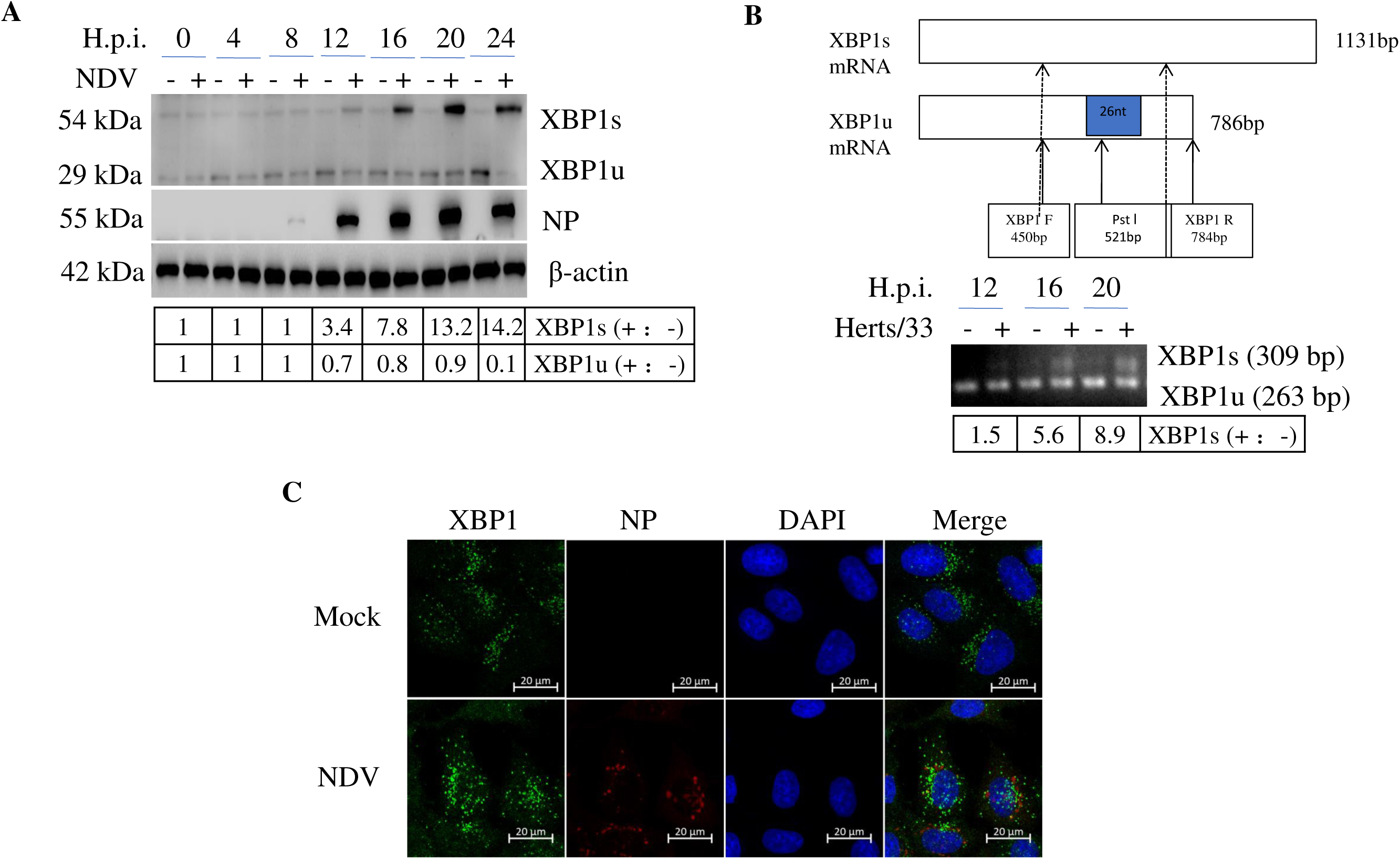
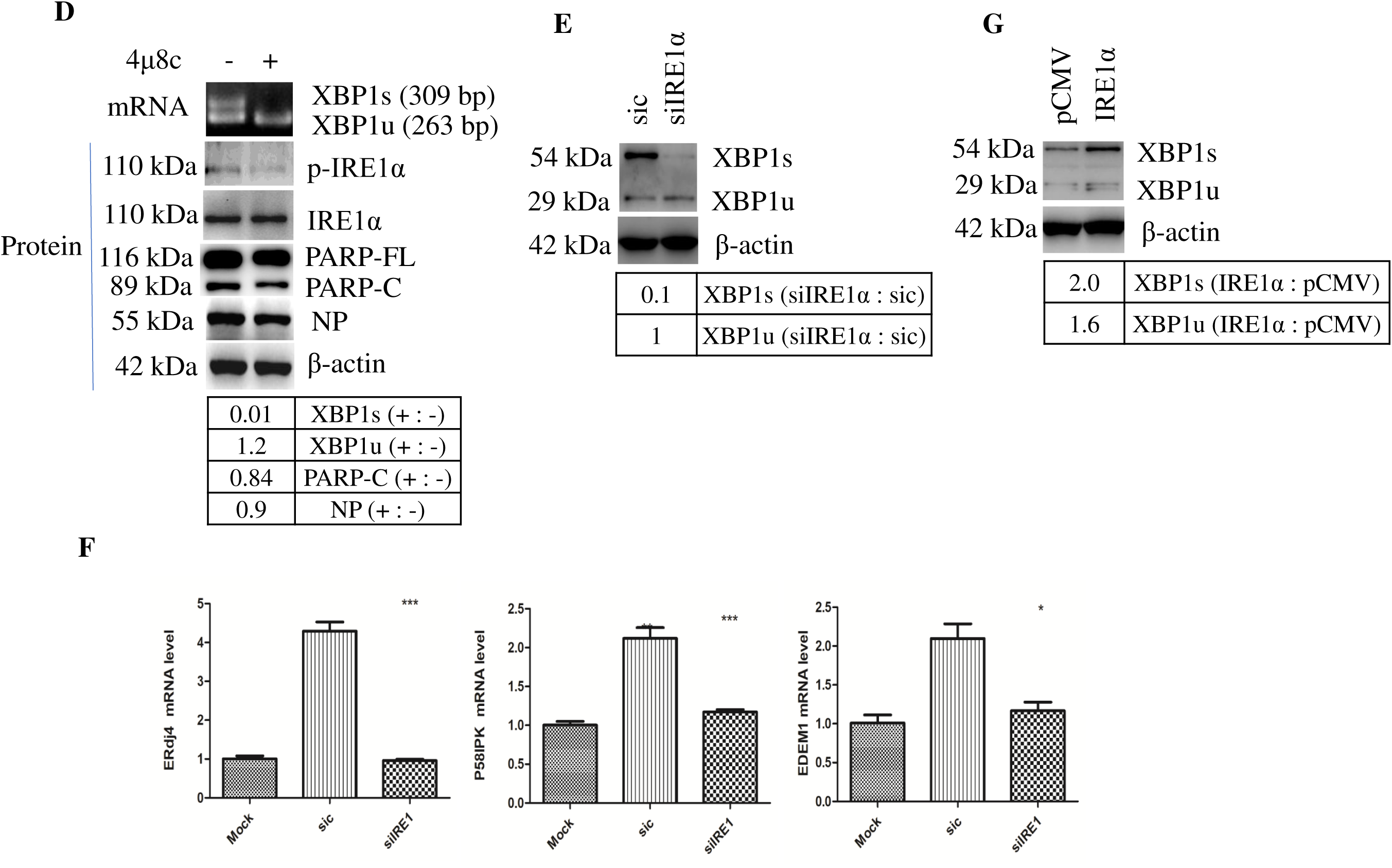
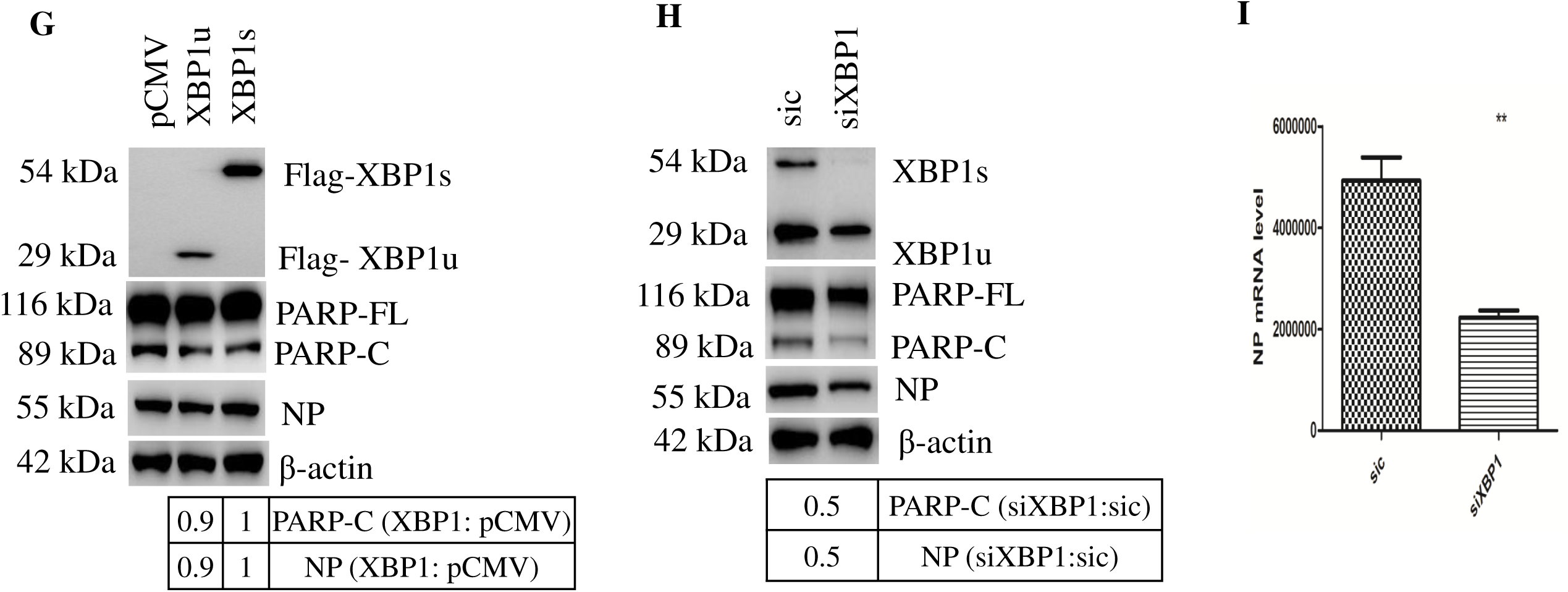
Splice of XBP1 by IRE1α promotes apoptosis and ERAD, and facilitates NDV proliferation. (A-B) NDV infection leads to XBP1 mRNA splicing and produces XBP1s. HeLa cells were infected with NDV or mock-infected, harvested at indicated time points, and analyzed with Western blotting (A) or RT-PCR (B) to detect the spliced form of XBP1. The intensity of XBP1u and XBP1s bands was normalized to β-actin and shown as fold change of NDV (+ : -) (A). RT-PCR was performed with XBP1 specific primers and the products were digested with *Pst* I. XBP1u products were 72 bp and 263 bp, XBP1s product was 309 bp. The 309 bp of XBP1s and 263 bp of XBP1u were shown in Fig. B. The intensity of XBP1s bands was determined and shown as fold change of NDV (+ : -). (C) The nuclear translocation of XBP1s during NDV infection. HeLa cells were infected with NDV or mock-infected, and subjected to immunofluorescence at 16 h.p.i. to detect XBP1 and NP (D) Inhibition of IRE1 RNase activity by 4μ8c blocks XBP1 mRNA splicing. HeLa cells were infected with NDV, treated with DMSO or 25 μM IRE1α RNase inhibitor 4μ8c, and subjected to RT-PCR or Western blotting analysis. RT-PCR was performed with XBP1 specific primers and the products were digested with *Pst* I. The 309bp of XBP1s and 263 bp of XBP1u were shown in upper panel. The intensity of XBP1s and XBP1u bands was determined and shown as fold change of 4μ8c (+ : -). The intensity of PARPHOSPHO-C and NP bands was normalized to PARPHOSPHO-FL or β-actin, and shown as fold change of 4μ8c (+ : -). (E-F) Knock down of IRE1α reduces XBP1 splicing, decreases chaperones and ERAD components expression. HeLa cells were transfected with silRElα or sic, followed with NDV infection for l6 h. Cells were analyzed with Western blotting to check the XBP1 splicing (E), or subjected to semi-quantitative real time RT-PCR to detect the mRNA level of IRE1α, p58^IPK^, ERdj4 and EDEMl (F). The intensity of XBP1s and XBP1u bands was determined by Image J software and normalized to the band intensity of β-actin, and shown as fold change of siIRE1α : sic (E). (G) Overexpression of IRE1α promotes XBP1 splicing. HeLa cells were transfected with pCMV-IRE1α or pCMV, followed with NDV infection. The cells were analyzed with Western blotting at l6 h.p.i. using XBP1 antibody. The intensity of XBP1s and XBP1u bands was normalized to β-actin and shown as fold change of IRE1α : pCMV. (G) Overexpression XBP1u slightly reduces apoptosis and virus proliferation. HeLa cells were transfected with pCMV-XBP1u, pCMV-XBP1s or pCMV, and infected with NDV. At l6 h.p.i., cell lysates were blotted with the primary antibodies against Flag, PARP, NP, and β-actin. The intensity of PARPHOSPHO-C and NP bands was normalized to PARPHOSPHO-FL or β-actin, and shown as fold change of XBP1: pCMV. (H-I) Knock down of XBP1 reduces apoptosis and virus proliferation. HeLa cells were transfected with siXBP1 or sic, and infected with NDV. At 16 h.p.i., cell lysates were analyzed with Western blotting using the indicated antibodies (H), or subjected to semi-quantitative real time RT-PCR to check the mRNA level of NP (I). The intensity of PARPHOSPHO-C and NP bands was normalized to PARPHOSPHO-FL or β-actin, and shown as fold change of siXBP1:sic.

IRE1α is responsible for splicing of XBP1 (27). We next examined the effect of IRE1α on XBP1 splicing during NDV infection. After NDV infection, IRE1α RNase activity was inhibited by 4μ8c, which specifically binds to the lysine residue in the ribonuclease catalytic pocket. DMSO treatment was included in a parallel experiment as control group. Cells were harvested at 16 h.p.i. and subjected to RT-PCR and Western blot. RT-PCR results showed that 4μ8c treatment markedly suppressed NDV-induced splicing of XBP1 mRNA, compared to that in DMSO treated cells (Fig. 5D). It was noted that the RNase inhibitor treatment only slightly reduced the PARP cleavage and NP protein synthesis (Fig. 5D), indicating the IRE1α RNase activity may not crucial for apoptosis and virus proliferation. To further confirm the role of IRE1α on XBP1 splicing, IRE1α was either knocked down or overexpressed, followed with NDV infection. As shown in Fig. 5E, compared with control group, knock down of IRE1α reduced the XBP1s protein to undetectable level; meanwhile, overexpression of IRE1α produced 2-fold of XBP1s protein (Fig. 5G). Accordingly, knock down of IRE1α reduced NDV-induced transcription of the ER chaperones and components of ERAD, including p58^IPK^, ERdj4 and EDEM1 genes, as evidenced by the semi-quantitative real time RT-PCR (Fig. 5F). Collectively, above results reveal that during NDV infection, IRE1α mediates the splicing of XBP1 mRNA, produces XBP1s protein as nuclear transcription factor, and initiates the transcription of ER chaperones and ERAD components.

### XBP1 is essential for efficient NDV replication

In order to maintain the homeostasis of the ER under stress, XBP1s induces the expression of ER chaperones and ERAD components, thereby enhancing the capacity of productive folding and degradation mechanism (12, 66). It is also reported that XBP1u and XBP1s is involved in IBV induced apoptosis (27). XBP1-dificient cells were resistant to apoptosis induced by vesicular stomatitis virus (VSV) and herpes simplex virus (HSV) infection (67). These reports indicate that XBP1 is involved in cell fate determination during virus infection. To study the role of XBP1u and XBP1s in NDV-induced apoptosis, we first adopted the overexpression approach. The coding sequence of XBP1u or XBP1s was inserted into pCMV vector respectively, with Flag tag at C-terminus. HeLa cells were transfected with construct XBP1u, XBP1s, or pCMV vector, followed with NDV infection. Using anti-Flag antibody, the expression of both proteins was clearly detectable (Fig. 5G). Compared with the vector control, in cells transfected with XBP1u, slightly lower level of NDV NP (0.9-fold) and NDV-induced PARP cleavage (0.9-fold) could be detected. In contrast, in cells transfected with XBP1s, the level NDV NP and NDV-induced PARP cleavage was similar to that in vector control (Fig. 5G). To further investigate the function of XBP1 in NDV-induced apoptosis and virus proliferation, we used siRNA to specifically knock down XBP1 in HeLa cells, followed with NDV infection. As expected, the expression of XBP1s was successfully knocked down by siXBP1, and XBP1u was moderately decreased (Fig. 5H). Interestingly, transfection of siXBP1 resulted in 0.5-fold decrease of NP synthesis, compared to those in sic transfected cells (Fig. 5H).

In consistence, NP mRNA was significantly decreased in siXBP1 transfected cells (Fig. 5I). Accordingly, PARP cleavage was decreased by 0.5-fold in siXBP1 transfected cells. Although overexpression of XBP1u or XBP1s has no significant effect on virus proliferation and virus induced apoptosis, the knock down experiment demonstrates that XBP1 is necessary for efficient NDV replication and NDV-induced apoptosis. However, it is difficult to attribute the observed phenotype to individual isoforms as siXBP1 targets both XBP1u and XBP1s.

### NDV infection activates pro-apoptotic JNK via IRE1α and NF-κB

In addition to mediating XBP1 mRNA splicing, IRE1α also recruits TRAF2 and ASK1, subsequently activating MKK4/7 and JNK (68). JNK promotes apoptosis either by directly regulating the apoptotic proteins activity or activating the transcription factor for pro-apoptotic protein (69). We have shown that JNK was phosphorylated at late stage of NDV infection (Fig. 3A), and CHOP promotes this activation (Fig. 3B, 3C). We next asked whether IRE1α was involved in NDV-induced JNK activation. HeLa cells were transfected with siIRE1α or sic before being infected with NDV, and the phosphorylation level of JNK was examined by Western blotting. As shown in Fig. 6A, knock down of IRE1α greatly reduced the phosphorylation of JNK by 0.2-fold, compared with that in sic control cells. In contrast, in cells transfected with plasmid encoding IRE1α, the NDV-induced phosphorylation of JNK was greatly increased by 2.2-fold, compared to that in vector transfected cells (Fig. 6B). Taken together, these data demonstrate that IRE1α promotes JNK phosphorylation in NDV-infected cells.

**Figure 6.**
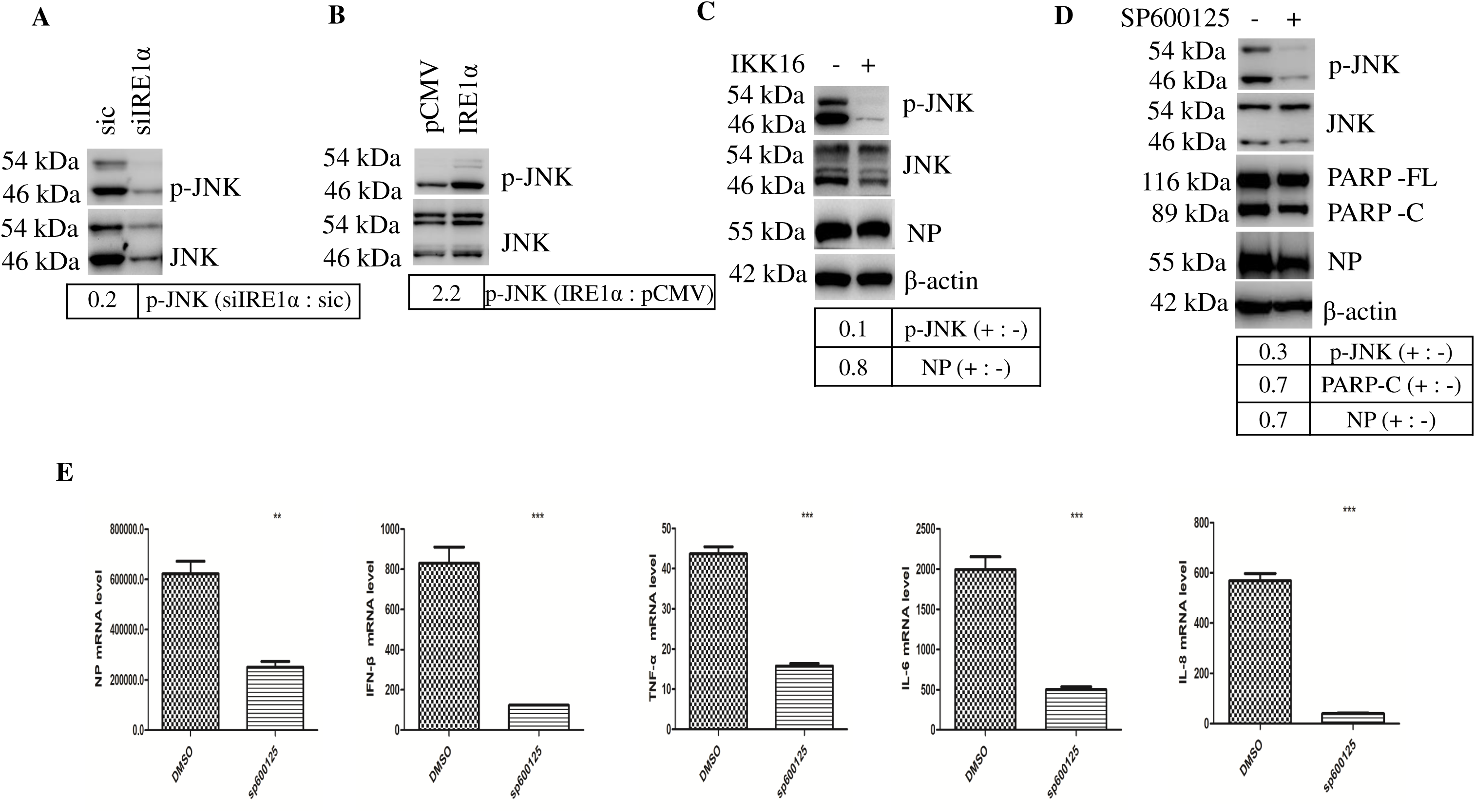
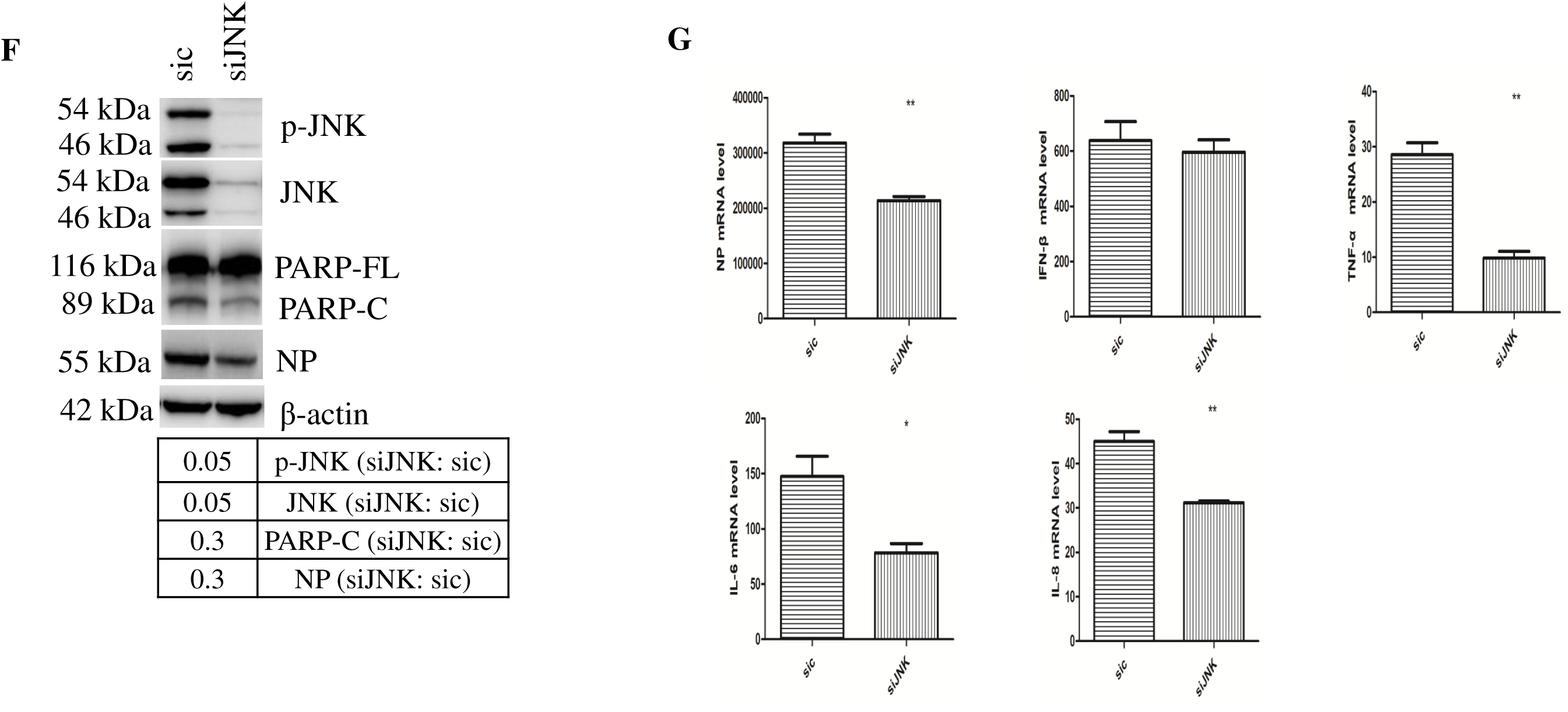
NDV infection activates pro-apoptotic and pro-inflammatory JNK signaling cascade via IRE1α and NF-κB. (A) Knock down of IRE1α decreases JNK signaling. HeLa cells were transfected with siIRE1α or sic, followed with NDV infection. At 16 h.p.i., cells were analyzed with Western blotting to check the phospho-JNK and JNK. The intensity of phospho-JNK bands was normalized to total JNK, and shown as fold change of silRElα : sic. (B) Overexpression of IRE1α promotes the activation of JNK signaling. HeLa cells were transfected with pCMV-IRE1α or pCMV, followed with NDV infection. AT 16 h.p.i., cells were analyzed with Western blotting to check the phospho-JNK and JNK. The intensity of phospho-JNK bands was normalized to total JNK and shown as fold change of IRE1α: pCMV. (C) Pharmacologic inhibition of NF-κB signaling suppresses JNK activation. HeLa cells were infected with NDV, and incubated with DMSO or 5 μM IKK16. At 16 h.p.i., the levels of phospho-JNK, JNK, NP, and β-actin were analyzed with Western blotting. The intensity of phospho-JNK and NP bands was normalized to total JNK or β-actin, and shown as fold change of IKK16 (+ : -). (D-E) Pharmacological inhibition of JNK activity by SP600125 protects cells from apoptosis and reduces NDV replication. HeLa cells were mock-infected or infected with NDV, followed by treatment with DMSO or 7.5 μM JNK inhibitor SP600125. The protein level of phospho-JNK, JNK, PARP, NP, and β-actin were analyzed with Western blotting. The intensity of phospho-JNK, PARPHOSPHO-C, and NP bands was normalized to total JNK, PARPHOSPHO-FL, or β-actin respectively, and shown as fold change of SP600125 (+ : -) (D). The mRNA levels of NP, IFN-β, TNF-α, IL-6, and IL-8 were determined with semi-quantitative RT-PCR using specific primers (E). (F-G) Knock down of JNK reduces apoptosis and virus proliferation. HeLa cells were transfected with siJNK or sic, followed with NDV infection. At 16 h.p.i., the levels of phospho-JNK, JNK, PARP, NP were analyzed with Western blotting using indicated antibodies. The intensity of phospho-JNK, PARPHOSPHO-C, and NP bands was normalized to total JNK, PARPHOSPHO-FL, or β-actin respectively, and shown as fold change of siJNK: sic (F). The mRNA levels of NP, IFN-β, TNF-α, IL-6, and IL-8 were determined with semi-quantitative RT-PCR using specific primers (G).

**Figure 7.**
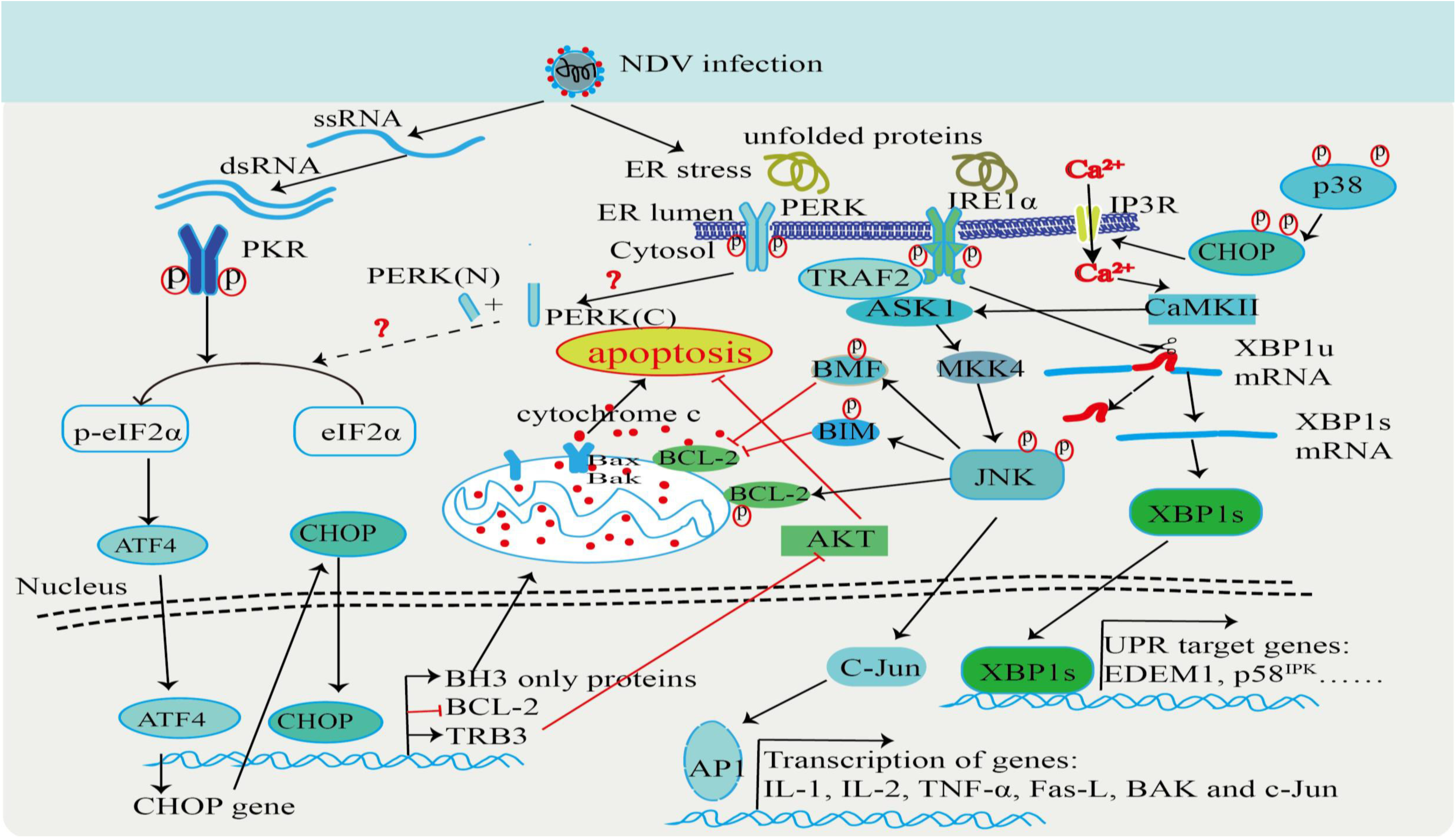
Working model of UPR associated apoptosis during NDV infection. NDV infection produces dsRNA, activates PKR and phosphorylates eIF2α, induces the expression of CHOP. CHOP promotes apoptosis by reducing the expression of anti-apoptotic protein BCL-2 and MCL-1, stimulating JNK and p38 signaling cascades, and inhibiting the pro-survival AKT signaling. Meanwhile, NDV infection results in ER stress and activates IRE1α-XBP1/JNK pathway. Both XBP1 and JNK signaling cascades promote apoptosis and benefit virus proliferation.

Previously, we reported that NDV infection activates NF-κB and induces TNF-α expression (44). TNF-α promotes apoptosis not only by activation of caspase 8 and NF-κB, but also by activation of JNK (70). We then checked whether NF-κB signaling also mediates the activation of JNK. IKKβ inhibitor IKK16 (5 μM) was incubated with NDV-infected cells to block the activation of NF-κB, and the phosphorylation level of JNK was checked. As shown in Fig. 6C, treatment with IKK16 did not change the expression of NDV NP protein; however, it reduced the phospho-JNK to minimum level (0.1-fold), compared to that in the control cells. This result suggests that JNK is not only activated by IRE1α, but also stimulated via NF-κB-TNF-α signaling. Thus, activation of JNK is controlled by multiple signaling during NDV infection.

To explore the role of JNK in NDV-induced apoptosis, JNK kinase activity was inhibited by SP600125 (7.5 μM) after NDV infection. Western blotting results showed that SP600125 treatment really inhibited the phosphorylation of JNK by 0.3-fold, and decreased NDV NP expression by 0.7-fold (Fig. 6D). Accordingly, in SP600125 treated cells, NP mRNA was also greatly decreased, compared to that in DMSO-treated cells (Fig. 6E). Meanwhile, inhibition of JNK kinase activity reduced the cleavage of PARP by 0.7-fold (Fig. 6D). Moreover, the transcription of death ligand TNF-α and cytokines IFN-β, IL-6, IL-8 was markedly suppressed in SP600125 treated cells, as evidenced by semi-quantitative real time RT-PCR (Fig. 6E). To validate above results, specifically knock down of JNK by siRNA was performed. As shown in Fig. 6F, JNK was successfully depleted by siJNK (0.05-fold), which significantly reduced the level of viral NP expression (0.3-fold), compared to that in sic control cells. PARP cleavage was also greatly decreased by 0.3-fold (Fig. 6F). Accordingly, NP, TNF-α, IL-6 and IL-8 mRNA was also suppressed by in JNK depletion cells (Fig. 6G). Collectively, above results demonstrate that activation of JNK promotes virus proliferation and virus-induced apoptosis/inflammation.

## DISCUSSION

During virus infection, many viral proteins are synthesized by ER-associated ribosome and transported into ER lumen for proper folding or post-translational modification. This leads to an overwhelming load of unfolded or misfolded proteins in ER lumen. Then, chaperone Bip binds to these unfolded/misfolded proteins and releases ER stress sensors PERK, ATF6, IRE1α, triggering UPR, marked as protein translation shut down, activation of transcription factors (ATF4, ATF6, XBP1s), expression of ER chaperones and ERAD. UPR determines cell fate to survival or death (71, 72). Many viruses have evolved mechanisms to manipulate host UPR signaling to help viral replication. For instance, dengue virus triggers IRE1α-XBP1 pathway to protect cells from virus induced cytopathic effects (73); Hepatitis C virus protein NS4B activates IRE1α to protect the infected cells from apoptosis, facilitating the development of chronic infection and hepatocellular carcinoma (67); Reovirus induces the phosphorylation of eIF2α and expression of ATF4, which activates the integrated stress response and promotes survival of stressed cells, benefiting virus replication (74); Classical swine fever virus (CSFV) activates IRE1-XBP1-GRP78 signal and maintains ER homeostasis, to promote its replication (75); Herpes simplex virus 1 (HSV-1) can flee from cellular responses that are likely detrimental to viral replication via suppressing the IRE1-XBP1 branch by tegument protein UL41 (76). Whether similar mechanisms are applied to NDV infection remains to be investigated

As an acute infection pathogen and oncolytic reagent, NDV induces apoptosis as a major hallmark in host cells and several tumor cell lines. However, whether UPR is involved in NDV-induced apoptosis has not been well characterized. Previous studies have shown that NDV infection activates PKR and promotes phosphorylation of eIF2α, resulting in preferential translation of ATF4, which enters into the nucleus and promotes the transcription of CHOP (45, 77). CHOP could suppress the expression of BCL-2 to release its sequestration pro-apoptotic proteins, BAX (78). CHOP also inhibits the activation of AKT, an anti-apoptosis kinase, by inducing the expression of TRB3 (79). Here, we find that NDV infection triggers the expression and nuclear translocation of CHOP via PKR-eIF2α signaling. Exogenous expression of CHOP promotes apoptosis by reducing the level of anti-apoptotic protein BCL-2 and MCl-1, while knock down of CHOP increases the level of these pro-survival proteins. MAPKs are canonical signaling pathways crosstalk with ER stress responses (80, 81). Indeed, NDV infection activates all three MAPKs: JNK, p38, ERK1/2. Meanwhile, CHOP promotes the NDV-induced pro-apoptotic JNK/p38 signaling cascades. It has been reported that CHOP indirectly promotes ER Ca^2+^ release, results in the activation of Ca^2+^ /calmodulin-dependent protein kinase II (CaMKII), subsequently promoting apoptosis through mitochondrial membrane potential loss or activation of ASK1-MKK4-JNK signaling cascade (82). Also, CHOP is also activated by p38-dependent phosphorylation (83, 84). The suppression of pro-survival AKT signaling by CHOP might be due to the expression of TRB3 (85). Thus, the NDV infection induced CHOP promotes apoptosis via regulation of BCl-2 family proteins, MAPK signaling, and AKT signaling. In addition to the pro-apoptotic role, CHOP is also essential for NDV proliferation.

IRE1α is a highly conserved ER stress sensor, which can be found organisms from yeast to mammals. Under ER stress, IRE1α is activated and splices XBP1u mRNA into XBP1s, which is subsequently translated into an active transcription factor (17). IRE1α also activates JNK to promote apoptosis (26). Therefore, IRE1α is involved in determination of cell fate (86). Previous studies have shown that IRE1α is activated by various virus infections, and viruses have different mechanisms to regulate IRE1α, XBP1, and JNK to facilitate their own replication. Hepatitis B virus, Influenza A virus, Japanese encephalitis virus, and Flavivirus activate IRE1-XBP1 branch, but Hepatitis C virus and Rotavirus suppress this pathway (87–92). Activation of IRE1α is helpful for the efficient replication of influenza A virus (87). The trans-activator protein VP16 of herpes simplex virus (HSV) can activate JNK pathway, which regulates the cell cycle, to promote successful virus replication (93). IRE1α protects cells from infectious bronchitis virus (IBV) induced apoptosis, which required both its kinase and RNase activities. The splicing of XBP1 mRNA by IRE1α convert XBP1 from a pro-apoptotic XBP1u protein to a pro-survival XBP1s protein (27). However, in a recent report, it was demonstrated that XBP1 deficiency confers resistance to intrinsic apoptosis by activation of IRE1α and decrease of miR-125a abundance, and results in increased virus infection (67). Thus, IRE1α and XBP1 may play either pro-apoptotic or anti-apoptotic role by different virus infection. In this study, we found that IRE1α was activated during NDV infection, and controlled XBP1 splicing and JNK activation. Exogenous expression of IRE1α sensitized cells to NDV induced apoptosis and enhanced the virus yield; while knock down of IRE1α protected cell from apoptosis and decreased virus yield. Consistent with these results, knock down of XBP1 protected cell from apoptosis and reduced virus yield. Both pharmacological inhibition of JNK and depletion of JNK by siRNA knock down reduced cell death and virus proliferation. Thus, the activation of UPR branch IRE1α-XBP1/JNK plays a pro-apoptotic role and helps NDV proliferation.

To facilitate shedding and dissemination of progeny viruses, some viruses take advantage of inducing apoptosis (94). NDV can specifically kill tumor cells by inducing apoptosis, then, this provides a promising therapeutic target for human tumors. Our current study demonstrates that NDV infection promotes apoptosis via inducing the expression of CHOP and activation of IRE1α-XBP1s/JNK, and the induction of these UPR branches or apoptosis is helpful for NDV proliferation. The full understanding of the involvement of these UPR branches in NDV replication process appears to be complicated. Probably, the expression of ER quality control proteins, which are controlled by IRE1α-XBP1 pathway, could promote virus replication by enhancing the viral proteins process. Another possibly is that the XBP1s could stimulate the phospholipid biosynthesis and ER expansion (95), providing the lipid that is necessary for the enveloped virus particle assembly. NDV-induced apoptosis may also help virus release. Meanwhile, apoptosis may avoid stimulating the anti-viral innate immune responses or inflammation in un-infected neighbor cells, in favor of next round infection. This study provides comprehensive insight into the mechanisms of ER stress induced apoptosis during NDV infection.

## ACKNOWLEDGEMENTS

This study was supported by National Natural Science foundation of China (Grant No. 31772724), Natural Science foundation of Shanghai (Grant No. 15ZR1449600), China Ministry of Science and Technology (Grant No. 2017YFD0500802), Elite Youth Program of Chinese Academy of Agricultural Sciences (Grant No. 20170260401), and National Natural Science foundation of China (Grant No. 31530074).

